# Single-cell analyses identify tobacco smoke exposure-associated, dysfunctional CD16^+^ CD8 T cells with high cytolytic potential in peripheral blood

**DOI:** 10.1101/783126

**Authors:** Suzanne N. Martos, Michelle R. Campbell, Oswaldo A. Lozoya, Brian D. Bennett, Isabel J.B. Thompson, Ma Wan, Gary S. Pittman, Douglas A. Bell

**Author notes:** These authors contributed equally (Suzanne N. Martos, Michelle R. Campbell).

## Abstract

Tobacco smoke exposure has been found to impact immune response, leukocyte subtypes, DNA methylation, and gene expression in human whole blood. Analysis with single cell technologies will resolve smoking associated (sub)population compositions, gene expression differences, and identification of rare subtypes masked by bulk fraction data. To characterize smoking-related gene expression changes in primary immune cells, we performed single-cell RNA sequencing (scRNAseq) on >45,000 human peripheral blood mononuclear cells (PBMCs) from smokers (n=4) and nonsmokers (n=4). Major cell type population frequencies showed strong correlation between scRNAseq and mass cytometry. Transcriptomes revealed an altered subpopulation of Natural Killer (NK)-like T lymphocytes in smokers, which expressed elevated levels of *FCGR3A* (gene encoding CD16) compared to other CD8 T cell subpopulations. Relatively rare in nonsmokers (median: 1.8%), the transcriptionally unique subset of CD8 T cells comprised 7.3% of PBMCs in smokers. Mass cytometry confirmed a significant increase (p = 0.03) in the frequency of CD16+ CD8 T cells in smokers. The majority of CD16+ CD8 T cells were CD45RA positive, indicating an effector memory re-expressing CD45RA T cell (T_EMRA_) phenotype. We expect that cigarette smoke alters CD8 T cell composition by shifting CD8 T cells toward differentiated functional states. Pseudotemporal ordering of CD8 T cell clusters revealed that smokers’ cells were biased toward later pseudotimes, and characterization of established markers in CD8 T cell subsets indicates a higher frequency of terminally differentiated cells in smokers than in nonsmokers, which corresponded with a lower frequency in naïve CD8 T cells. Consistent with an end-stage T_EMRA_ phenotype, *FCGR3A*-expressing CD8 T cells were inferred as the most differentiated cluster by pseudotime analysis and expressed markers linked to senescence. Examination of differentially expressed genes in other PBMCs uncovered additional senescence-associated genes in CD4 T cells, NKT cells, NK cells, and monocytes. We also observed elevated T_regs_, inducers of T cell senescence, in smokers. Taken together, our results suggest smoking-induced, senescence-associated immune cell dysregulation contributes to smoking-mediated pathologies.

## INTRODUCTION

As a risk factor for human diseases, the global disease burden attributed to tobacco smoke exposure remains substantial. The World Health Organization (WHO) estimates approximately six million deaths per year from tobacco smoke exposure, resulting from both chronic and communicable diseases (Stampfli and Anderson, 2009; WHO, 2015). In smokers, a decline in immunity and increased risk of inflammatory diseases, such as atherosclerosis, argues that smoking-associated diseases are mediated by immune dysfunction. The development and progression of atherosclerotic lesions serves as an example of a complex immune-mediated pathology because T cells, monocytes, macrophages, dendritic cells (DCs), and B cells have all been reported to be involved (Hansson, 2005; Ilhan and Kalkanli, 2015). Refining smoking-associated changes in specific cells within immune populations will enhance our understanding of how dysfunctional immune subsets arise from exposure to tobacco smoke. This will facilitate prevention of diseases by identifying immune cells to target for clinical intervention.

In addition to DNA damage, smoking alters the epigenome and transcriptome of human blood leukocytes (Reynolds et al., 2017; Su et al., 2016; Wan et al., 2018). In Su et al. (2016), we demonstrated that many changes identified in isolated cell fractions, which correspond to major immune populations, were distinct from each other and whole blood. For example, *ITGAL*, which is expressed in T cells and involved in inflammation (Wang et al., 2014), had significantly decreased methylation in smokers’ T cells but not in whole blood or isolated cell fractions. It follows that bulk data from isolated fractions, which are comprised of multiple subtypes, would similarly mask meaningful changes, especially when differences arise in low frequency subsets. As such, interpretation of bulk genomic approaches is limited because changes could indicate altered distribution of cell (sub)populations or changes in expression within (sub)populations. The recent development of single cell methods provides the technology to resolve smoking-associated (sub)population composition changes, gene expression differences, and identification of rare subtypes obscured by bulk fraction data. Additionally, the multiparameter data allows us to concordantly study multiple cell types from the same individuals.

To identify cell (sub)populations affected by smoking and possibly connect observed immune cell changes with smoking-associated diseases, we characterized gene expression profiles and cell surface marker phenotypes from primary peripheral blood mononuclear cells (PBMCs) from four nonsmokers and four smokers by single cell RNA sequencing (scRNAseq) and mass cytometry. The combination of transcriptome profiling and immunophenotyping provides higher confidence in the validity of our findings than one single cell method alone. Major cell type population frequencies showed strong correlation between scRNAseq and mass cytometry. In addition to resolving PBMCs into major immune cell types, we used single-cell transcriptome profiling to separate cell populations into multiple subsets according to differentiation, activation, or functional states. We discovered a rare population of CD16+ CD8 T cells that was increased in smokers and exhibited NK-like transcriptional programs. Pseudotime analysis and examination of canonical markers revealed that these NK-like CD8 T cells likely represent a terminally differentiated state. Not unique to CD8 T cells, other immune populations also displayed genes characteristic of senescence in smokers.

By discovering an altered abundance of a rare population, single cell methodologies revealed a novel immune target that can be isolated and explored for connections between smoking and chronic diseases. Combined with increased (pre-)senescent CD8 T cells, elevated regulatory T cells (T_regs_) and induction of senescence-linked genes in multiple cell types provide evidence that smokers show signs of premature aging of their immune systems. The potential immune function defects and inflammatory subsets demonstrated here mirror characteristics of pathologies commonly found in smokers. Further studies of smoking-associated dysregulation of immune transcriptional programs and candidate dysfunctional T cells linked to accelerated aging of the immune system, uncovered here, will lead to mechanistic insights to advance disease prevention strategies for smoking-mediated pathologies.

## RESULTS

### scRNAseq and Mass Cytometry Profiling of Human Peripheral Blood Immune Cells in Smokers and Nonsmokers

We set out to characterize the effects of cigarette smoke on immune cells in peripheral blood using single-cell approaches to determine whether smoking-associated gene expression changes observed within major immune cell populations resulted from altered abundance of specific, identifiable cell subsets. We performed scRNAseq and mass cytometry, in parallel, on cryopreserved peripheral blood samples from eight donors (Figure 1A). The samples were obtained from nonsmokers (n = 4) and smokers (n = 4) with no previous history of atherosclerosis, chronic obstructive pulmonary disease (COPD), or lung cancer. We used serum cotinine, a metabolite of nicotine and established biomarker of recent cigarette smoke exposure (Florescu et al., 2009), to confirm smoking status of donors. Smokers used for single-cell analyses had serum cotinine levels ranging from 240 – 511 ng/ml; all nonsmokers had serum cotinine levels below 2 ng/ml. Nonsmoking donors were matched to smoking donors based on gender and race. Donors’ ages ranged from 31 – 56 and were not significantly different between smokers and nonsmokers (p = 0.23). Demographic and smoking information for each individual used in this study is listed in Table S1. We obtained single-cell mRNA data from 45,049 cells and surface protein expression data for 26 markers from 990,748 cells.

**Figure 1.**
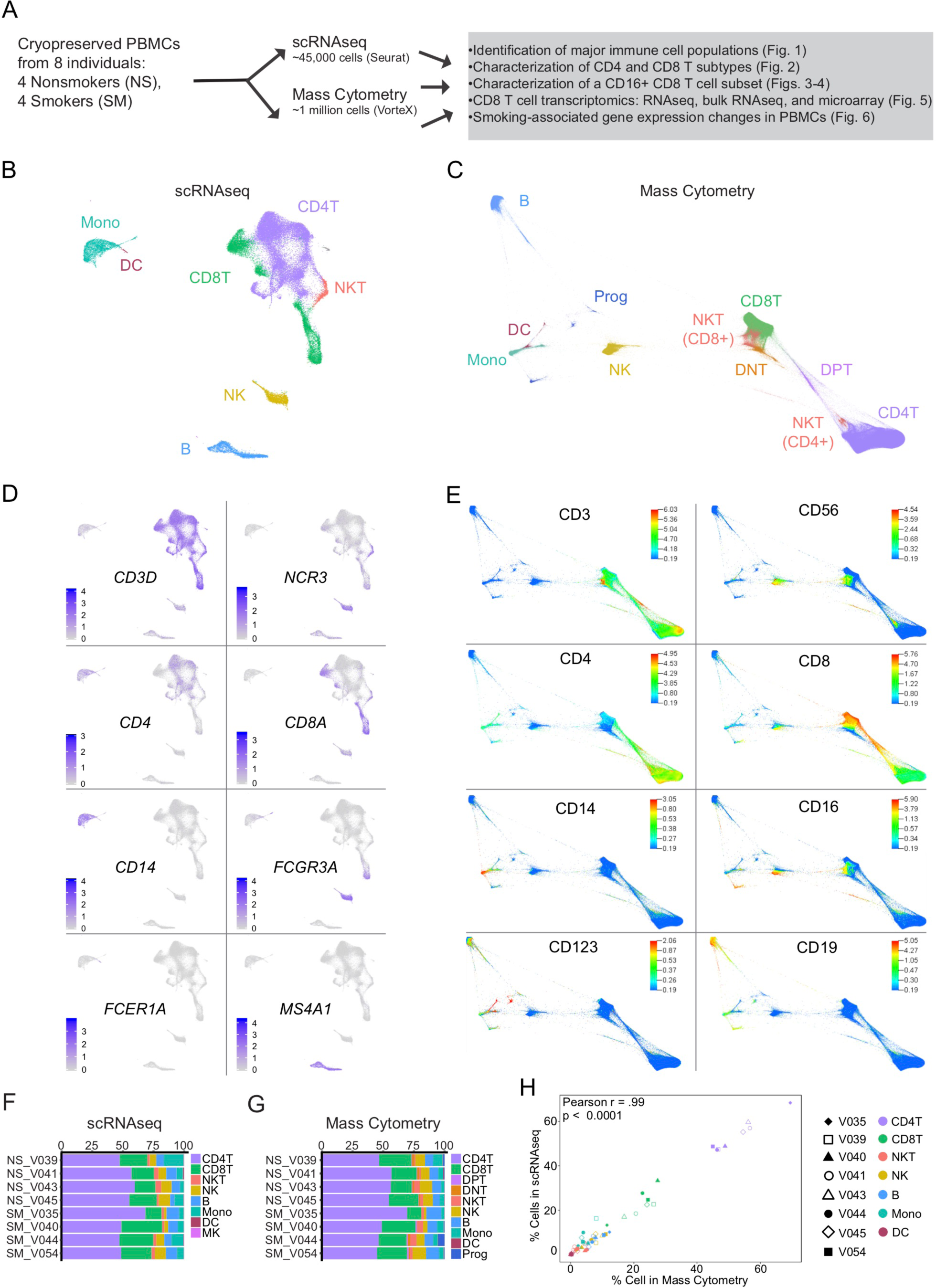
Profiling of single cell RNA sequencing (scRNAseq) and mass cytometry of human peripheral blood mononuclear cells (PBMCs) from smokers and nonsmokers. (A) Overview of experimental design. Cryopreserved PBMCs from smokers and nonsmokers were thawed for scRNAseq and mass cytometry, in parallel. (B) Uniform Manifold Approximation and Projection (UMAP) plot of single cell RNA sequencing (scRNAseq) showing ∼45,000 human peripheral blood mononuclear cells (PBMCs) colored by major cell types. (C) Force Directed Layout (FDL) of mass cytometry showing ∼1 million live cells colored by major cell types. (D) scRNAseq displaying canonical gene expression markers for major cell types: CD8T cells (*CD3D, CD8A*), CD4T cells (*CD3D, CD4*) Natural Killer T (NKT) cells (*CD3D, NCR3*), Natural Killer (NK) cells (*NCR3*), monocytes (*CD14, FCGR3A* [CD16]), dendritic cells (DCs; *FCER1A*) and B cells (*MS4A1*). (E) Mass cytometry displaying cell surface protein expression for major cell types: CD8T cells (CD3, CD8a), CD4T cells (CD3, CD4), NKT cells (CD3, CD56), NK cells (CD56), monocytes (CD14, CD16), DCs (CD123) and B cells (CD19). (F-G) Major cell type population frequency distributions shown by individual donor (Nonsmokers, NS; Smokers, SM) for scRNAseq (F) and mass cytometry (G), colored by cell type. (H) Major cell type population frequencies showed strong correlation between scRNAseq and mass cytometry (Pearson r = .99, r2 = 0.98, p<0.0001). Shapes represent paired individuals, nonsmokers (unfilled) and smokers (filled), are colored by cell type.

For each single-cell approach, we assigned cells to common immune populations based on mRNA (scRNAseq) or surface protein expression (mass cytometry) of well-characterized markers (Figures 1B-1E and Table S2). For scRNAseq, we used Seurat (Butler et al., 2018; Stuart et al., 2019) to anchor datasets across donors and implement shared nearest neighbors (SNN) clustering (see Methods). We then used Model-based Analysis of Single-cell Transcriptomics (MAST; (Finak et al., 2015)) to identify positive and negative marker genes for each cluster and combined cells into major immune populations based on expression of marker genes (Table S2). Cells in clusters expressing *CD3D* as a positive marker were designated as T cells (Figure 1D). T cells were further classified into CD4 T cells, CD8 T cells, or NKT cells based on expression of *CD4, CD8A*, or *NCR3* (Figure 1D. NK cells were identified based on *CD3D* as a negative marker combined with expression of *NKG7, GNLY, GZMB, PRF1*, and *NCR3* as positive markers (Figure 1D and Table S2). Monocytes were positive for *LYZ* and either *CD14* or *FCGR3A* (gene encoding CD16 protein), characteristic of classical or nonclassical monocytes (Figure 1D). Dendritic cells were similar to monocytes but could be distinguished by expression of *FCER1A* (Figure 1D). B cells were defined by *MS4A1* (gene encoding CD20 protein) (Figure 1D).

In parallel, 250,000 PBMCs from each donor were assessed by mass cytometry using an immunophenotyping panel (see Methods). Viable, single-cell events were manually gated to remove normalization beads, doublets and dead cells using Cytobank ((Kotecha et al., 2010); Figure S1A) and imported into the VorteX Clustering Environment using default parameter recommendations (Samusik et al., 2016). Using weighted k-nearest neighbor clustering, an elbow point validation was performed to determine the optimal clustering k value which was then used to create a Force-Directed Layout (FDL) graph using the X-shift algorithm (see methods). 122 PBMC cell cluster identities (IDs) were determined from the eight donors representing 983,848 cells (Figure S1B) in which the cells could also be visualized by smoking status (Figure S1C). Cell surface protein expression profiles were used to identify the cell populations (Figures 1C and 1E). Cells identified as T cells displayed CD3 (Figure 1E), which could then be classified with CD4 and CD8 (Figure 1E) as double negative (DNT), double positive (DPT), CD4 T, or CD8 T cells (Figure 1C). NKT cells were identified by CD3 and CD56 with either CD4 or CD8 protein expression markers. Monocytes were identified by protein expression of CD14 and/or CD16 and dendritic cells had CD123 above background levels (Figure 1E). B cells were positive for CD19 (Figure 1E). NK cells were positive for CD56 but negative for CD3 (Figure 1E).

To determine how well the scRNAseq and mass cytometry corresponded with each other, we examined the individual donor contribution in each cluster for each cell type. Cells colored by individual donors are shown for scRNAseq (Figure S1D) and mass cytometry (Figure S1E). Cell type frequencies were calculated based on cluster identification and plotted to compare frequency distributions among individuals (Figures 1F and 1G). For both methods, all major populations—CD4T, CD8T, NKT, B, Monocyte, and DC—were identified in all donors. We then compared the frequency of major populations in PBMCs by smoking status for scRNAseq and mass cytometry using a Mann Whitney U test. We observed no differences in the overall frequency of major cell types between smokers and nonsmokers by either scRNAseq or mass cytometry (Figures S1F and S1G). Comparing the percentage of cells in the major immune populations among individuals for scRNAseq and mass cytometry showed significant strong correlation (Pearson r = 0.99, r2 = 0.98, p<0.0001) between methods (Figure 1H).

### scRNAseq Reveals Increased T_regs_ and Altered Composition of the CD8 T Cell Population Between Smokers and Nonsmokers

In addition to dividing PBMCs into major immune cell types, single-cell transcriptome profiling can be used to separate cell populations into multiple subsets according to differentiation, activation, or functional states. Based on gene expression patterns, we clustered peripheral blood cells into thirty-one immune cell clusters and one erythroid contaminant cluster, labeled based on abundance from 0 (most abundant) through 31 (Figure 2A and Table S2). We identified twelve CD4 T cell (0, 1, 3, 4, 6, 10, 13, 14, 17, 18, 20, and 27), seven CD8 T cell (2, 8, 11, 15, 19, 21, and 24), three NK cell (7, 22, and 25), four monocyte (5, 23, 26, and 30), and two B cell (9 and 12) clusters. NKTs (16), DCs (28), and MKs (31) were each contained by a single cluster. Clusters are referred to as major immune cell type, followed by original cluster ID (e.g., CD4T-0). To determine whether smoking altered the subtype distribution within the major cell populations, we compared the abundance of cells among clusters for each major cell type that separated into more than one cluster. Cells colored by smoking status are shown in Figure 2B. We did not observe any subset frequency shifts in B cells, monocytes, or NK cells (Figures S2A-S2F).

**Figure 2.**
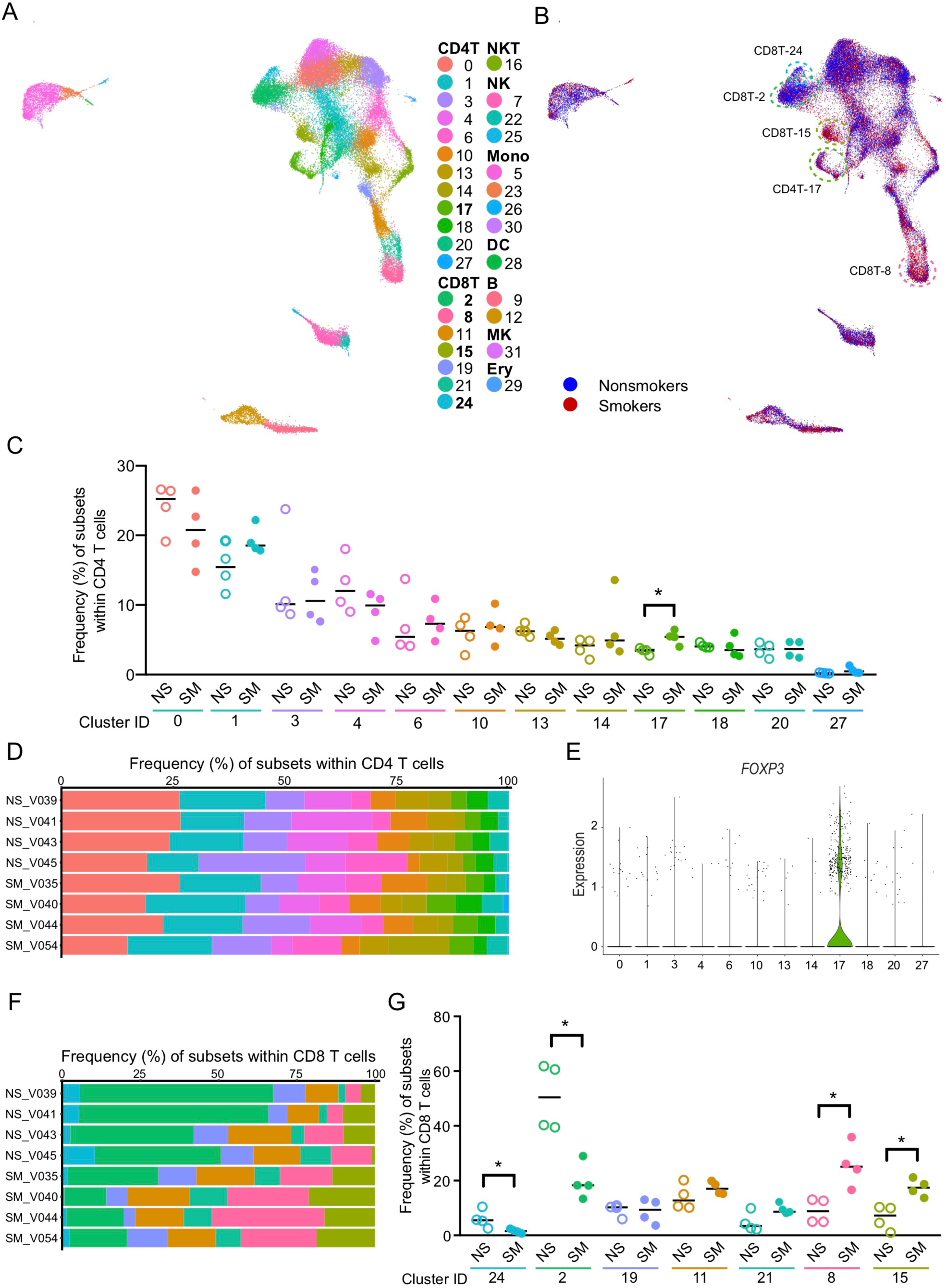
An increase in CD4 T_regs_ and an altered composition in the CD8 T cell population is observed between smokers (SM) and nonsmokers (NS) (A) scRNAseq UMAP colored by cluster ID number. (B) scRNAseq UMAP colored by smoking status (nonsmokers blue, smokers red). Clusters CD8T-24 and CD8T-2 had more cells from nonsmokers and CD4T-17, CD8T-15 and CD8T-8 showed more cells from smokers. Dotted circles indicate cluster ID. (C-D) Frequency of subsets within CD4 T cells. Smokers (filled) had a significant increase in cluster CD4T-17 compared to nonsmokers (unfilled). Bar = median, *p < 0.05 by Mann-Whitney U test (C). Individual donor distribution of CD4 T cell subset*s* (D). (E) Violin plot of *FOXP3* gene expression within the CD4 T subsets. (F-G) Frequency of subsets within CD8 T cells. Individual donor distribution of CD8 T cell subsets (F). Smokers (filled) had significant decreases in clusters CD8T-24 and CD8T-2 and increases in clusters CD8T-8 and CD8T-15 compared to nonsmokers (unfilled). Bar = median, *p < 0.05 by Mann-Whitney U test (G).

For 11 of 12 CD4 T cell subsets, frequency was not significantly altered by smoking (Figures 2B and 2C). Although donors exhibited interindividual variation in percent of each CD4 T cell subset, smoking status did not appear to have a considerable impact on the distribution of CD4 T cells (Figure 2D). Only one cluster, CD4T-17, was higher in smokers than in nonsmokers (p < 0.05; Figure 2C). This cluster had relatively low abundance among CD4 T cell subsets: median 3.5% in nonsmokers and 5.4% in smokers. We characterized CD4T-17 cells as regulatory T cells (T_regs_) based on elevated *FOXP3* and *IL2RA* (gene encoding CD25) compared to other PBMCs (Figure 2E and Table S2). No other CD4T clusters showed either *FOXP3* and *IL2RA* as strong positive markers.

In contrast to CD4 T cells, variation among donors in CD8 T cells appeared to depend on smoking status as illustrated by distinct differences of the dominant population(s) in nonsmoking and smoking donor CD8 T cells (Figure 2E). Smokers had lower proportions of two CD8 T cell clusters (CD8T-24 and CD8T-2) and higher proportions of two CD8 T cell clusters (CD8T-8 and CD8T-15) compared to nonsmokers (p < 0.05; Figure 2F). We did not observe significant differences in proportions of the remaining three CD8 T cell clusters (CD8T-19, CD8T-11, CD8T-21).

### Altered Distribution of CD8 T Cell Subsets Indicates Shift from Naïve to Differentiated CD8 T Cell States in Smokers

Smoking had substantial effects on the composition of CD8 T cells. Because of this, we further analyzed each of the seven CD8 T clusters to identify gene expression patterns that made cells in each cluster distinct. We used MAST to identify positive and negative markers for each CD8 T cluster relative to other CD8 T cells (see Methods). We found 71, 223, 46, 144, 71, 739, and 105 positive and 86, 978, 10, 68, 59, 214, and 52 negative markers (q < 0.05 in both smokers’ and nonsmokers’ cells) for CD8T-24, CD8T-2, CD8T-19, CD8T-11, CD8T-21, CD8T-8, and CD8T-15 clusters, respectively (Table S3). We next examined marker gene lists for genes associated with T cell differentiation and function to distinguish between CD8 T subsets. Several genes frequently used to classify CD8 T subsets varied among CD8 T cell clusters (Figures 3A and 3B). Elevated *CCR7, SELL*, and *IL7R* combined with low *CCL5* indicate that clusters CD8T-24 and CD8T-2 represent naïve CD8 T cells. High levels of *IL7R, SELL*, and *FOS* (associated with proliferation of activated T cells (Martins et al., 2008; Shaulian and Karin, 2002)) suggest that cluster CD8T-15 consists of cells exhibiting characteristics of long-lived memory cells, such as central memory T cells (T_CM_). The reduced levels of *CCR7, SELL*, and *IL7R* and elevated *CCL5* and *KLRG1* in CD8T-11, CD8T-21, and CD8T-8 are indicative of later T cell differentiation stages (e.g., T_EM_). The lack of *CD27* expression and decrease in *FOS* in CD8T-21 and CD8T-8 suggest highly differentiated T_EM_ cells (i.e. T_EMRA_). *ZEB2*, associated with terminal differentiation states (Scott and Omilusik, 2019), was detected in approximately one-third of CD8T-8 cells (35.1% in smokers and 28.2% in nonsmokers).

**Figure 3.**
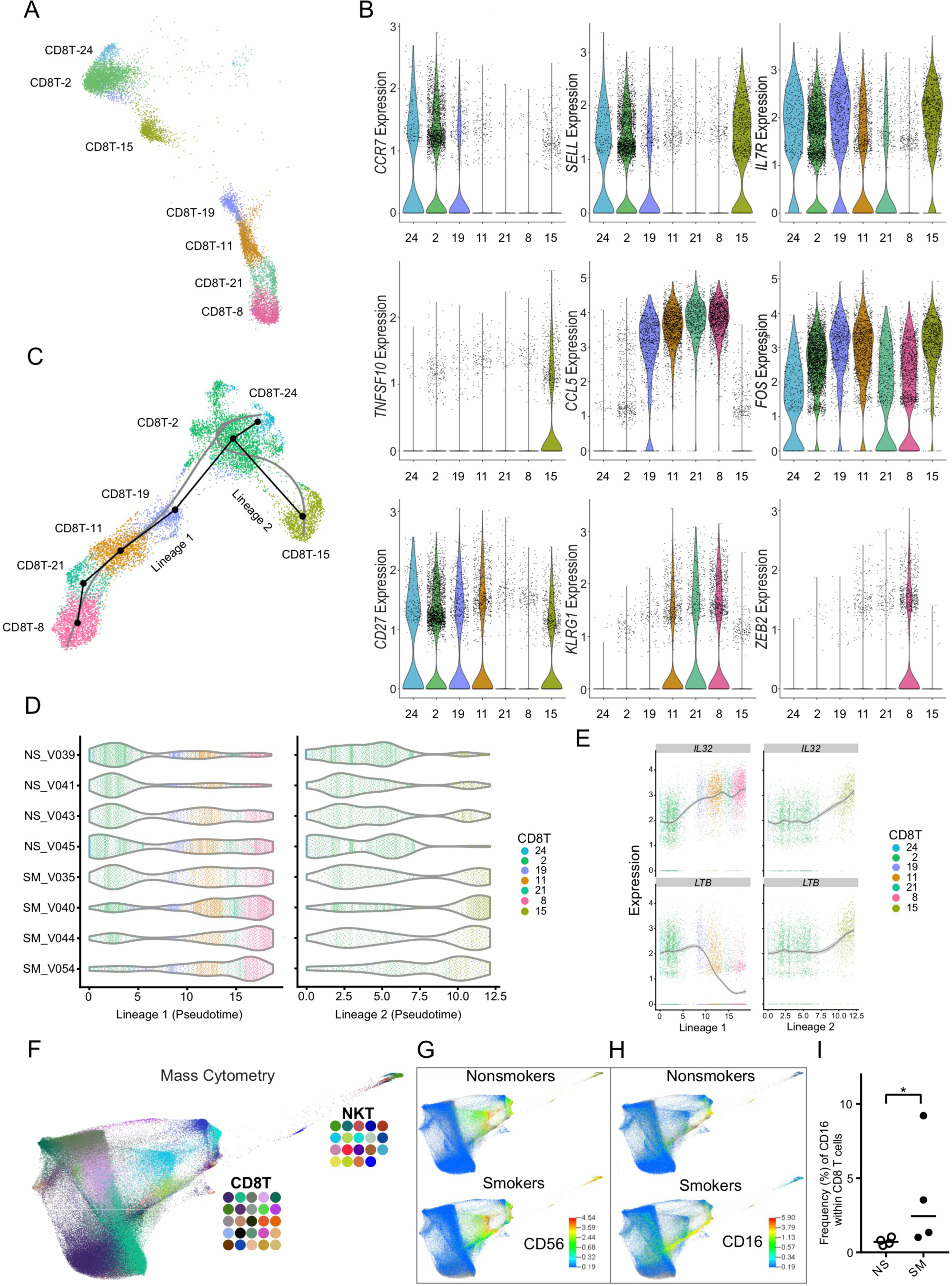
CD8 T cell subsets shift from naïve to differentiated CD8 T cells states. (A) scRNAseq UMAP of the seven CD8 T cell subsets colored by cluster ID number. (B) Violin plots showing expression of genes selected to characterize CD8 T subsets. Color indicates cluster ID. (C) Trajectory inference (pseudotime analyses) ordered CD8 T cell clusters into two lineages, which originate from cluster CD8T-24 and terminate at either cluster CD8T-8 (lineage 1) or CD8T-15 (lineage2). Black line shows lineage tree and the gray line shows the simultaneous principle curves. (D) Violin plots show pseudotime for CD8 T cells by individual nonsmokers (NS) and smokers (SM) in lineage one (left) and lineage two (right). (E) Pseudotemporal trajectory of CD8 T cell differentiation in *IL32* (top) and *LTB* (bottom) for lineage one (left) and lineage two (right). (F-I) Mass cytometry confirms elevated proportion of CD16+ CD8 T cells in smokers. (F) FDL of CD8 T and NKT cells colored by cluster ID. (G-H) Cell surface marker intensity FDL plots of nonsmokers (top) and smokers (bottom) for NK marker CD56 (G) and CD16 (H). (I) Frequency of CD16+ cells increases within CD8 T cells in smokers (filled) compared to nonsmokers (unfilled). Bar = median, *p < 0.05 by one-tail Mann-Whitney U test.

Several clusters exhibited intermediate expression of differentiation-state genes. To organize CD8 T cell clusters by their likely differentiation trajectories, we used the Slingshot algorithm (Street et al., 2018) to perform pseudotemporal analysis. Lineage inference ordered CD8 T cells into two lineages, which originate from cluster CD8T-24 and terminate at either cluster CD8T-8 or CD8T-15 (Figure 3C). Lineage one was mostly comprised of cells from CD8T-24, CD8T-2, CD8T-19, CD8T-8, CD8T-21, and CD8T-8, with minimal cells from CD8T-15. Lineage two was mostly comprised of cells from CD8T-24, CD8T-2, and CD8T-15, with minimal cells from CD8T-19. Based on the altered composition of CD8 T cell subsets—lower proportions of CD8T-2 and CD8T-24 and higher proportions of CD8T-8 and CD8T-15 cells (Figures 2B, 2F, and 2G)—we propose that tobacco smoke exposure alters CD8 T cell composition by shifting CD8 T cells toward differentiated states. Smokers’ cells were biased toward later pseudotimes in both lineages (Figure 3D), demonstrating smokers’ CD8 T cells are skewed toward differentiated and nonsmokers’ CD8 T cells are skewed toward naïve states.

After ordering cells by pseudotime, we identified temporally associated genes for each lineage. Only 10 genes within the top 100 temporally expressed genes (Figures S3A and S3B) were shared between the lineages. *IL32*, which is induced in activated T cells (Goda et al., 2006), increased in both (Figures 3E). Similarly, most shared genes exhibited the same direction of change over pseudotime in both lineages. In contrast, *LTB*, a TNF superfamily ligand (Ware, 2005), decreased over pseudotime in lineage one and increased over pseudotime in lineage two. In general, temporally expressed genes for CD8 T cell lineages were consistent with effector memory (lineage one) and central memory (lineage two) differentiation. For example, in lineage one, *CCL5* and *NKG7* increased, while *SELL* and *IL7R* decreased over the differentiation trajectory (Figures S3A and S3C). *CMC1* demonstrated a nonlinear association in lineage one, as it peaked in CD8T-21 cells and then decreased through CD8T-8 cells (Figures S3A and S3C).

The terminal cluster in the effector memory trajectory, CD8T-8, shared many features with the penultimate cluster, CD8T-21; however, CD8T-8 increased in smokers but CD8T-21 did not. Since the low expression of *CD27* and *CCR7* and elevated expression of *KLRG1* in both clusters would classify these cells as highly differentiated CD8 T cells (i.e. T_EMRA_-like), we sought to find markers within CD8T-8 that did not occur in CD8T-21 to detect distinguishing features of this cluster. Examination of exhaustion markers—*TOX, PDCD1* (gene encoding PD-1), *CTLA4*, and *HAVCR2* (gene encoding TIM3) did not distinguish the clusters. Whereas smokers’ CD8T-8 cells had elevated expression of *TOX* (logFC = 0.22, q = 1.44 x 10^−9^), nonsmokers’ CD8T-8 cells did not, and no other exhaustion markers were significantly elevated in CD8T-8 cells (Table S3). We next examined senescence-associated genes *KLRG1* and *B3GAT1* (gene encoding CD57). While *KRLG1* was considered a positive marker for CD8T-21 (smokers: logFC = 0.45, q = 8.11 x 10^−04^; nonsmokers: logFC = 0.92, q = 2.23 x 10^−22^) and CD8T-8 cells (smokers: logFC = 0.65, q = 6.29 x 10^−47^; nonsmokers: logFC = 0.89, q = 1.94 x 10^−40^), *B3GAT1* was unique to CD8T-8 cells (smokers: logFC = 0.14, q = 3.83 x 10^−14^; nonsmokers: logFC = 0.07, q = 0.022). We also found that two genes reported as having smoking-associated methylation changes, *GFI1* and *PRSS23* (Joehanes et al., 2016), showed elevated expression in CD8T-8 cells (*GFI1* smokers: logFC = 0.14, q = 5.58 x 10^−7^; *GFI1* nonsmokers: logFC = 0.15, q = 0.046; *PRSS23* smokers: logFC = 0.74, q = 1.5 x 10^−143^; *PRSS23* nonsmokers: logFC = 0.63, q = 7.7 x 10^−60^). Surprisingly, *FCGR3A*, which is commonly found on NK cells and nonclassical monocytes, was identified as a strong positive marker of CD8T-8 cells in smokers (logFC = 1.13, q = 1.17 x 10^−171^) and nonsmokers (logFC = 1.20, q = 5.17 x 10^−119^; Figure 1D and Table S3).

### Mass Cytometry Confirms Elevated Proportion of CD16+ CD8 T Cells in Smokers

Relatively rare in nonsmokers (median: 1.8%), the *FCGR3A*-expressing CD8 T cell cluster (CD8T-8) comprised 7.3% of PBMCs in smokers. Reported as a low-frequency subset (∼ 2% of PBMCs in healthy adults), CD16^+^ CD8 T cells have been described previously (Bjorkstrom et al., 2008; Clemenceau et al., 2008; Clemenceau et al., 2011). Based on these reports and similarities of cells in the CD8T-8 cluster with cells in other effector memory CD8 T cell clusters (CD8T-11 and CD8T-21), we sought to ascertain whether smokers had increased levels of CD16^+^ CD8 T cells that expressed surface proteins for CD3, but not CD56. That is, confirm an increase in CD16 expression within CD8 T cells that are not NKT cells. To show that smokers had increased surface protein expression of CD16 within their CD8 T cells, we first ran the X-shift algorithm on all CD8 T and NKT cells (Figures 3F-H). After visualizing the FDL colored by cluster IDs (Figure 3F) we examined the protein expression intensities for CD56 and CD16 (Figures 3G and 3H) for smokers and nonsmokers. We did not see any differences between smokers’ and nonsmokers’ CD8 T cells for CD56 (Figure 3G), but smokers showed an increase in the proportion of CD8 T cells expressing CD16 compared to nonsmokers (Figure 3H). In order to determine the frequency of CD16^+^ CD8 T cells in smokers and nonsmokers, a biaxial plot for CD3 and CD56 was created for manual gating (Figure S3D). CD3^+^CD56^-^ negative cells were then gated by CD4 and CD8 to obtain single positive CD8 T cells (CD3^+^CD56^-^CD8^+^CD4^-^, Figure S3D), which were then used to determine the frequency of CD16^+^ CD8 T cells (CD3^+^CD56^-^CD8^+^CD4^-^CD16^+^, Figure S3D). Compared to nonsmokers, smokers had a significant increase in the frequency of CD16^+^ CD8 T cells (p = 0.03, Figure 3I) confirming that smokers had elevated proportions of CD16^+^ CD8 T cells.

In order to phenotype the CD16^+^ CD8 T cell subset, we gated CD3+ T cells by a CD45RA/CD45 biaxial plot to establish an accurate CD45RA^+^ gate that was then applied to the CD16^+^ CD8 T cells (Figure S3E). The majority of CD16^+^ CD8 T cells were positive for CD45RA in both smokers and nonsmokers (Figure S3F). Thus, this CD8 T subset is likely comprised of effector memory T cells re-expressing CD45RA (T_EMRA_) cells.

### *FCGR3A*(CD16)-expressing CD8 T Cells Exhibit Transcriptome Signatures Characteristic of a Natural Killer-like Phenotype

After confirming an increase in CD16^+^ CD8 T cells in smokers, we further examined how the transcriptomes of *FCGR3A*-expressing CD8 T cells differed from other CD8 T cells. In addition to *FCGR3A*, CD8T-8 cells exhibited elevated expression *NKG7, GNLY, FGFBP2, GZMB*, and *PRF1* (Tables S2 and S3). While these genes can be considered expression signatures typical of both cytotoxic T cells and NK cells, the presence of CD16 lead us to suspect that this subset might express additional genes indicative of NK-like attributes. To gain further insight into the functional relevance of gene expression profiles for CD8T-8 cells, we performed Gene Set Enrichment Analysis (GSEA). Consistent with an NK-like transcriptional program, LI_INDUCED_T_TO_NATURAL_KILLER_UP had a positive normalized enrichment score (NES = 1.88, FWER < 0.05) and GSE22886_NAIVE_TCELL_VS_NKCELL_UP had a negative NES (−2.13, FWER < 0.05). The LI_INDUCED_T_TO_NATURAL_KILLER_UP gene set encompasses expression patterns for T cells reprogrammed to have NK-like phenotypes: “induced T to NK” (iTNK) cells (Li et al., 2010). Here, we found elevated expression of 67 and 71 genes from the iTNK gene signature as positive markers (q < 0.05) of *FCGR3A*-expressing CD8 T cells in smokers and nonsmokers, respectively. Figure 4A shows the 25 highest ranked positive markers (based on q-value) for CD8T-8 cells within the iTNK gene set. In addition to positive enrichment for iTNK genes, CDT8-8 genes were negatively enriched (FWER < 0.05) for genes that are higher in naïve CD8 T cells relative to NK cells (GSE22886_NAIVE_TCELL_VS_NKCELL_UP). Smokers’ and nonsmokers’ CD8T-8 cells had significantly reduced expression (i.e. negative markers) of 55 and 45 genes within the naïve CD8 T vs NK gene set. Figure 4B shows the 25 highest ranked negative markers (based on p-value) for CD8T-8 cells within the naïve CD8 T vs NK gene set.

**Figure 4.**
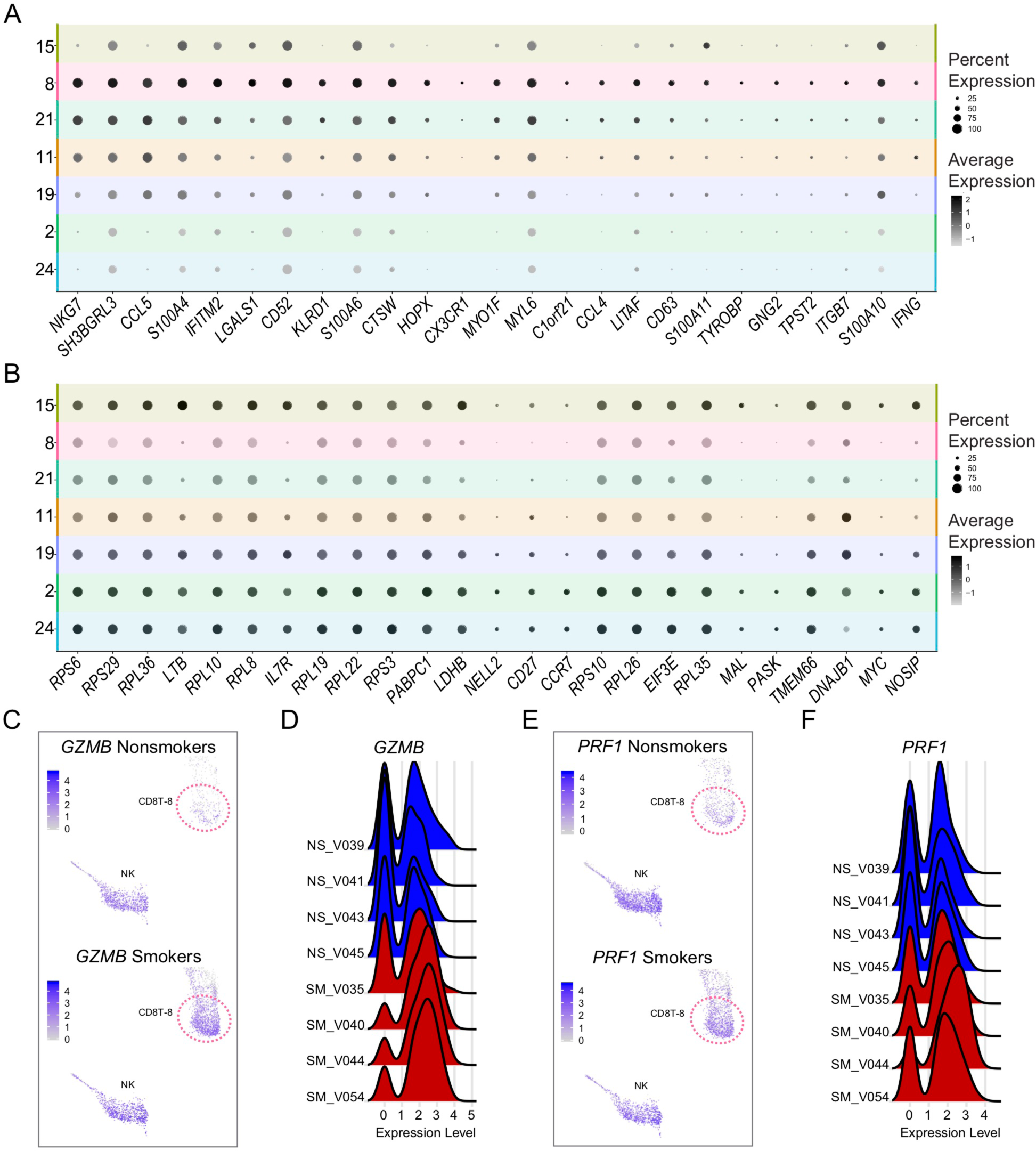
Transcriptome characterization of CD16+ CDS T cells. (A-B) Gene Set Enrichment Analysis (GSEA) of gene expression in CD8T-8 cells compared to cells in other CD8 T clusters revealed NK-associated transcriptome characteristics. CD8T-8 cells were positively enriched (FWER <0.05) for gene signatures of T cells induced to have NK-like phenotypes; split dot plot shows 25 genes within the T induced to NK gene set (A). CD8T-8 cells were negatively enriched (FWER <0.05) for genes that are higher in na”fve CD8 T cells relative to NK cells; split dot plot shows 25 genes within the na”fve T vs NK gene set (B). Color intensity represents average per-cell expression (scaled shows standard deviation) and circle size indicates percentage of cells expressing each gene. (C-D) Cluster CD8T-8 cells show increased *GZMB* expression in smokers. (C) UMAP comparison of *GZMB* between nonsmokers’ (top) and smokers’ (bottom) cells. Cluster CD8T-8 is indicated by dotted circle. (D) Ridge plots displaying individual donor *GZMB* expression levels for CD8T-8 cells for nonsmokers (NS) and smokers (SM). (E-F) Cluster CD8T-8 cells have increased *PRF1* expression in smokers. (E) UMAP comparison of *PRF1* between nonsmokers’ (top) and smokers’ (bottom) cells. Cluster CD8T-8 is indicated by dotted circle. **(F)** Ridge plots displaying individual donor *PRF1* expression levels for CD8T-8 cells for nonsmokers (NS) and smokers (SM).

To examine how smokers’ “NK-like” CD8 T cells differed from those of nonsmokers, we compared average expression of genes from smokers’ CD8T-8 cells to nonsmokers’ CD8T-8 cells (see Methods). We found that 63 genes had increased and 74 genes had decreased average per cell expression in smokers compared to nonsmokers (q < 0.05; Table S4). Figures S4A and S4B show 25 genes with increased and decreased per cell expression, ordered by difference in percentage of cells expressing each gene between smokers’ and nonsmokers’ NK-like CD8 T cells. Although cellular mRNA levels for effector molecules granzyme B (encoded by *GZMB*) and Perforin (encoded by *PRF1*) exhibited interindividual variation, both increased in frequency of expression and average per cell expression in smokers compared to nonsmokers (Figures 4C – 4F, S4A, S4C and S4D).

### CD8 T Bulk Transcriptomes Reflect Differentiation-State Shifts Observed at Single-Cell Level

To assess the overall impact of smoking on CD8 T cells, we identified differentially expressed genes (DEGs) between smokers and nonsmokers by comparing the average per cell expression for all cells in the seven CD8 T clusters combined. Of 2163 genes evaluated in the pseudobulk analysis, we found that 1817 genes had higher expression and 344 genes had lower expression in smokers versus nonsmokers (q < 0.05; Table S5). To examine the interindividual variability in response to smoking, we performed hierarchical clustering using smoking scRNA-DEGs, which separated individual donors by smoking status (Figures 5A and S5A). To confirm altered CD8 T gene expression profiles in smokers, we used RNAseq and microarray on isolated CD8 T cells to examine differences in bulk RNA expression between smokers and nonsmokers. Isolated CD8 T cells for bulk RNAseq included the eight donors used in scRNAseq and seven additional donors (seven smokers, eight nonsmokers); samples obtained from the donors used in scRNAseq were from a previous visit (Table S1). We identified 1268 genes as differentially expressed; 692 increased and 576 decreased (q < 0.05; Figure 5B). With exception of F061, principal component analysis of bulk RNAseq data separated smokers from nonsmokers (Figure 5C). We also evaluated microarray data from isolated CD8 T cells from 19 donors (9 smokers and 10 nonsmokers). Isolated CD8T cells included 4 donors used in scRNAseq, 4 donors used in bulk RNAseq, and 11 additional donors (Table S1). We identified 51 genes as differentially expressed (see Methods); 46 increased and 5 decreased (Figure 5D).

**Figure 5.**
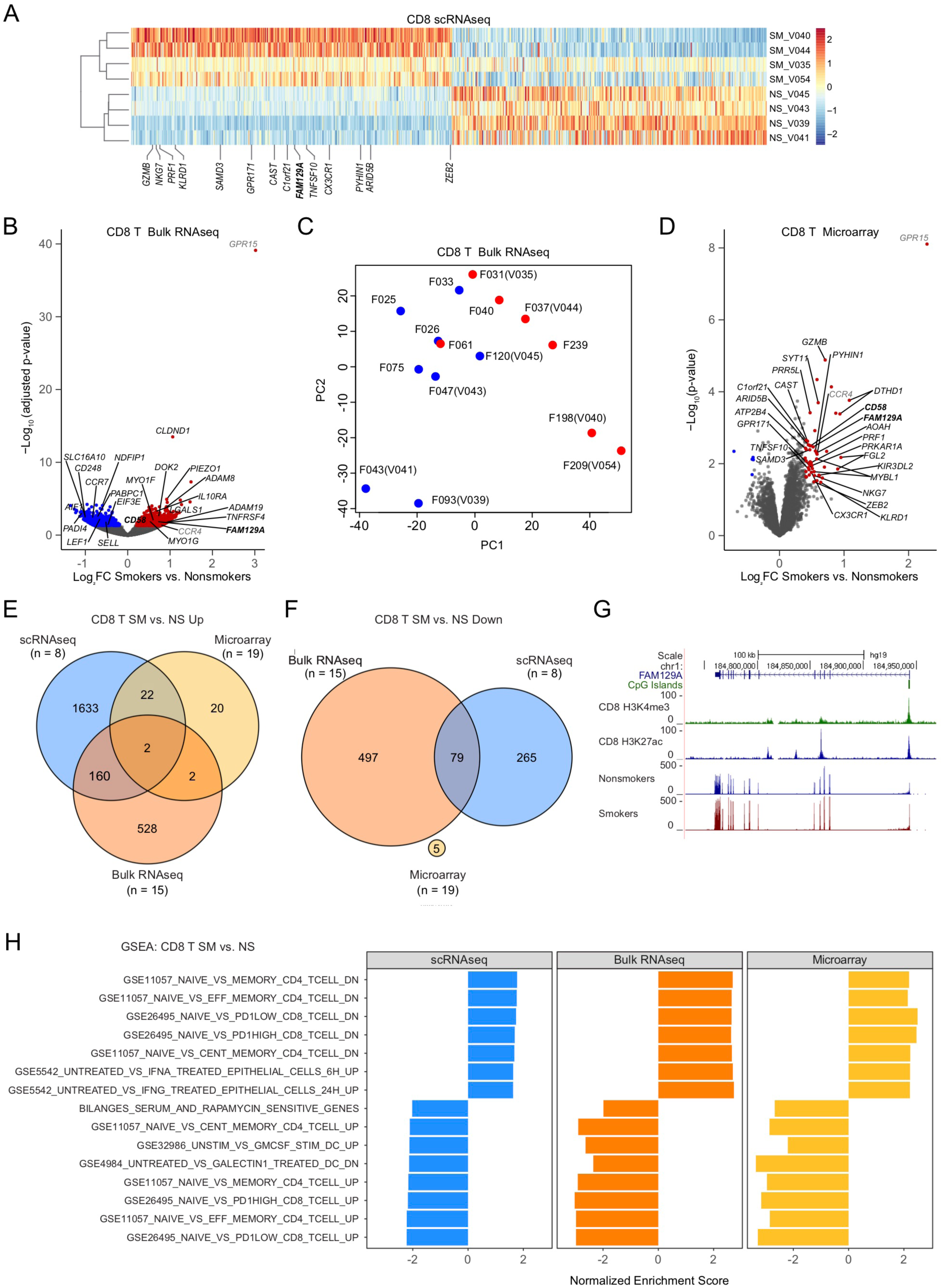
CD8 T bulk transcriptomes reflect differentiation-state shifts at single-cell level. (A) scRNAseq heatmap of the top 350 upregulated and downregulated differentially expressed genes (DEGs) between smokers and nonsmokers from the seven CD8 T cell clusters. Individual donors were separated by smoking status using smoking scRNA-DEGs for hierarchical clustering. Genes labeled were also found to be significantly upregulated in the CD8 T cell microarray results. (B) Volcano plot displaying bulk RNAseq expression of isolated CD8 T cells between smokers and nonsmokers. Genes with higher expression in smokers are colored red and genes with lower expression in smokers are colored blue. Gene names in black were altered in bulk RNAseq and scRNAseq. Gene names in gray were altered in bulk RNAseq and microarray. Gene names in bold were altered in microarray, bulk RNAseq and scRNAseq. (C) Principal component analysis (PCA) of bulk RNAseq data from an expanded group of donors. Isolated CD8 T cells from the same donors as included in scRNAseq and mass cytometry data were from a previous donor visit and are indicated with both an “F” and “V” patient code. See Table S1 for donor information. (D) Volcano plot displaying microarray expression of isolated CD8 T cells between smokers and nonsmokers, see Table S1 for details. Genes with higher expression in smokers are colored red and genes with lower expression in smokers are colored blue. Gene names in black were altered in microarray and scRNAseq. Gene names in gray were altered in microarray and bulk RNAseq. Gene names in bold were altered in microarray, bulk RNAseq and scRNAseq. (E-F) Venn diagrams comparing DEGs among scRNAseq, microarray and bulk RNAseq for upregulated (E) and downregulated genes (F) in smokers compared to nonsmokers. (G) Genome browser tracks of bulk RNAseq data for *FAM129A*, which was significantly increased by all three methods. (H) GSEA pathways for immunological signatures and chemical and genetic perturbations that were significantly enriched (FWER < 0.05) for scRNAseq, bulk RNAseq and microarray.

We next compared results from the three methods (pseudobulk scRNAseq, bulk RNAseq, and bulk microarray) used to identify smoking-associated DEGs in CD8 T cells. Bulk RNAseq confirmed 241 smoking DEGs from the scRNAseq, 162 with increased and 79 with decreased expression in smokers compared to nonsmokers (Figures 5B, 5E and 5F; Table S5). Microarray analysis confirmed 24 smoking DEGs that were identified in the scRNAseq with increased expression in smokers compared to nonsmokers (Table S5; Figures 5A, 5D and 5E). Two genes, *FAM129A* and *CD58*, were identified as increased in smokers’ CD8 T cells by all three methods (Figures 5A – 5B, 5D – 5E, 5G, and S5A; Table S5). Several genes found to be altered by at least two methods include *LGALS1, ADAM8, and CLDND1*, which were significantly increased in bulk RNAseq and scRNAseq data; *GPR15*, which was increased in the bulk RNAseq and microarray data; and *NDFIP1*, which was significantly decreased in the bulk RNAseq and scRNAseq data (Figures 5A, 5B, 5D, and S5B – S5E). *ITGAL*, a smoking methylation biomarker (Su et al., 2016), was only found to be significantly increased by scRNAseq (logFC = 0.25, q = 1.7 x 10^−26^), and was also elevated in NK-like subset (smokers: logFC = 0.45, q = 6.53 x 10^−29^; nonsmokers: logFC = 0.54, q = 1.50 x 10^−20^). Although each method identified smoking DEGs not found by other methods, we expect the overall gene expression changes observed for each method to represent a similar shift in the functional states of CD8 T cells. We used GSEA to determine whether CD8 T pseudobulk and bulk transcriptomes were enriched for similar functional annotations. There were seven positively and eight negatively enriched gene sets in common among pseudobulk scRNAseq, bulk RNAseq, and bulk microarray (FWER < 0.05; Figures 5H, 5SF, and 5SG). CD8 T cells were positively enriched for genes with higher expression in memory, central memory, effector memory, PD1 low (CD8 T effector memory), and PD1 high (CD8 T effector memory) T cells relative to naïve T cells (Figure 5H). CD8 T cells were negatively enriched for genes that have higher expression in naïve T cells relative to memory, central memory, effector memory, PD1 low (CD8 T effector memory), and PD1 high (CD8 T effector memory) T cells (Figure 5H). Therefore, GSEA of CD8 T smoking DEGs identified immunological signatures indicative of increased expression of genes associated with effector memory and central memory functions and decreased expression of genes associated with naïve T cells by all three methods.

### Smoking-associated Gene Expression Changes in PBMC Populations

Since we did not observe substantial changes in subset distribution for most of the major PBMC populations, we compared average per cell expression of genes for all cells within each cell type (CD4 T, NKT, NK, Monocyte, DC, and B) to identify smoking DEGs. For CD4 T cells, we found 1563 DEGs; 1278 showed increased expression and 285 showed decreased expression (Table S6). Hierarchical clustering of donors by CD4 T smoking DEGs clustered individuals by smoking status (Figures 6A). Analysis of bulk gene expression in isolated CD4 T cells by microarray identified two upregulated (*LRRN3* and *GPR171*) and one downregulated (*APBA2*) gene in common with the CD4 T pseudobulk analysis (Figures 6A and S6A). NKT cells had 89 smoking DEGs, 45 with increased and 44 with decreased expression, and NK cells had 238 smoking DEGs, 129 with increased and 109 with decreased expression (Table S6). Hierarchical clustering of DEGs separated donors by smoking status for NKT, but not for NK cells (Figures 6B and 6C). Microarray analysis of isolated CD56 cells, which contain NKT and NK cells, shared three upregulated genes and one downregulated gene with NKT cells and three upregulated and three downregulated genes with NK cells (Figures 6B, 6C, and S6B). *MX1*, which was increased in smokers’ CD56 cells, showed increased expression in both NKT and NK cells (Figures 6B and 6C). *KLRB1*, which was decreased in smokers’ CD56 cells, showed decreased expression in both NKT and NK cells (Figures 6B and 6C). For monocytes, we found 488 DEGs between smokers and nonsmokers by scRNA pseudobulk analysis (Figure 6D and Table S6). Of the 290 DEGs with higher expression in smokers’ monocytes, 15 were also shown to be increased in smokers’ isolated CD14 cells by microarray (Figures 6D and S6C). For DCs, we identified 21 smoking DEGs; 5 showed increased expression and 16 showed decreased expression (Table S6). Hierarchical clustering of DC smoking DEGs separated donors into two groups: three smokers and four nonsmokers with one smoker (Figure 6E). For B cells, we found 190 DEGs; 111 with increased and 79 with decreased expression in smokers (Table S6). Using microarray in isolated CD19 cells, we were able to confirm decreased expression of *HLA-DQA1* in smokers’ B cells compared to nonsmokers’ B cells (Figures 6F and S6D). Table 1 lists the biological relevance of smoking DEGs that were found in major PBMC populations by pseudobulk scRNAseq and confirmed by microarray.

**Table 1.** Smoking DEGs altered in both scRNAseq and microarray.

| Gene Name | DEG | Biological Process/Function/Pathway |
| --- | --- | --- |
| **CD8 T** |  |  |
| granzyme B | ↑ *GZMB* | PD1low (Duraismamy et al., 2011) |
| natural killer cell granule protein 7 | ↑ *NKG7* | PD1low, PD1high (Duraismamy et al., 2011) |
| perforin 1 | ↑ *PRF1* | PD1low, PD1high (Duraismamy et al., 2011) |
| killer cell lectin like receptor D1 | ↑ *KLRD1* | PD1low, PD1high (Duraismamy et al., 2011) |
| sterile alpha motif domain containing 3 | ↑ *SAMD3* | CD8 T tolerance (Schietinger et al., 2012; Uniken Venema et al., 2019) |
| G protein-coupled receptor 171 | ↑ *GPR171* | Negative regulation of myeloid differentiation (Rossi et al., 2013) |
| calpastatin | ↑ *CAST* | PD1low, PD1high (Duraismamy et al., 2011); Eff Memory, Cent Memory, (Abbas et al., 2009) |
| chromosome 1 open reading frame 21 | ↑ *C1orf21* | Cytotoxic T by scRNAseq in Crohn's Disease (Uniken Venema et al., 2019) |
| family with sequence similarity 129, member A | ↑ *FAM129A* | PD1low, PD1high (Duraismamy et al., 2011); Memory, Eff Memory, Cent Memory (Abbas et al., 2009) |
| TNF superfamily member 10 | ↑ *TNFSF10* | Marker for atherosclerosis plaque formation (Arcidiacono et al., 2018) |
| C-X3-C motif chemokine receptor 1 | ↑ *CX3CR1* | Chemokine Receptor Associated with terminally differentiated effector CD8 cells (Gerlach et al., 2016) |
| pyrin and HIN domain family member 1 | ↑ *PYHIN1* | PD1low, PD1high (Duraismamy et al., 2011); Memory (Abbas et al., 2009) |
| AT-rich interaction domain 5B | ↑ *ARID5B* | Cent Memory (Abbas et al., 2009) |
| zinc finger E-box binding homeobox 2 | ↑ *ZEB2* | PD1low, PD1high (Duraismamy et al., 2011) |
| CD58 molecule | ↑ *CD58* | PD1low, PD1high (Duraismamy et al., 2011); Memory, Eff Memory, Cent Memory (Abbas et al., 2009) |
| protein kinase cAMP-dependent type I regulatory subunit alpha | ↑ *PRKAR1A* | Enhance cytotoxicity of Effector T cells (Panya et al., 2018) |
| acyloxyacyl hydrolase | ↑ *AOAH* | Eff Memory Phenotype, increase in CLL CD3 T cells (Gothert et al., 2013) |
| proline rich 5 like | ↑ *PRR5L* | PD1low, PD1high (Duraismamy et al., 2011); Eff Memory, Cent Memory (Abbas et al., 2009) |
| MYB proto-oncogene like 1 | ↑ *MYBL1* | PD1low, PD1high (Duraismamy et al., 2011); Memory, Eff Memory, Cent Memory (Abbas et al., 2009) |
| ATPase plasma membrane Ca2+ transporting 4 | ↑ *ATP2B4* | PD1low, PD1high (Duraismamy et al., 2011); Memory, Eff Memory, Cent Memory (Abbas et al., 2009) |
| synaptotagmin 11 | ↑ *SYT11* | PD1low, PD1high (Duraismamy et al., 2011) |
| killer cell immunoglobulin like receptor, three Ig domains and long cytoplasmic tail 2 | ↑ *KIR3DL2<sup>a</sup>* | T cell aging (Chen et al., 2013) |
| death domain containing 1 | ↑ *DTHD1* | Unknown function |
| fibrinogen like 2 | ↑ *FGL2<sup>a</sup>* | Replicative senescence (Binet et al., 2009); procoagulant (Hu et al., 2016) |
| **CD4 T** |  |  |
| leucine rich repeat neuronal protein 3 | ↑ *LRRN3<sup>a</sup>* | Smoking methylation (Guida et al., 2015); T cell replicative senescence (Chou et al., 2013) |
| G protein-coupled receptor 171 | ↑ *GPR171* | Negative regulation of myeloid differentiation (Rossi et al., 2013) |
| amyloid beta precursor protein binding family A member 2 | ↓ *APBA2* | Smoking methylation (Chung et al., 2018) |
| **NKT** |  |  |
| prokineticin 2 | ↑ *PROK2* | Inflammatory response in smoking (Yun et al., 2017) and diabetes (Castelli et al., 2016) |
| MX dynamin like GTPase 1 | ↑ *MX1<sup>a</sup>* | TNF-α induced cellular senescence (Kandhaya-Pillai et al., 2017) |
| T cell receptor associated transmembrane adaptor 1 | ↑ *TRAT1* | Leukocyte activation (Taylor et al., 2017) |
| killer cell lectin like receptor B1 | ↓ *KLRB1<sup>a</sup>* | Downregulated in human aged CD4+ memory T cells (Chen et al., 2013) |
| **NK** |  |  |
| MX dynamin like GTPase 1 | ↑ *MX1<sup>a</sup>* | TNF-α induced cellular senescence (Kandhaya-Pillai et al., 2017) |
| dedicator of cytokinesis 5 | ↑ *DOCK5<sup>a</sup>* | Upregulated in CD57+ NK (Lopez-Verges et al., 2010) |
| chondroitin sulfate N-acetylgalactosaminyltransferase 1 | ↑ *CSGALNACT1* | Positively correlated with smoking (Charlesworth et al., 2010) |
| CD160 molecule | ↓ *CD160<sup>a</sup>* | Increased in exhausted T cells (Kasakovski et al., 2018) |
| X-C motif chemokine ligand 2 | ↓ *XCL2* | Overexpressed in lung cancer tumors, increases with prognosis (Zhou et al., 2016) |
| killer cell lectin like receptor B1 | ↓ *KLRB1<sup>a</sup>* | Downregulated in human aged CD4+ memory T cells (Chen et al., 2013) |
| **Monocytes** |  |  |
| phospholipid scramblase 1 | ↑ *PLSCR1* | Expression increased with cytokine treatment (Kodigepalli et al., 2015) |
| GTPase, IMAP family member 4 | ↑ *GIMAP4* | Regulates INF-gamma in CD4 T cell differentiation (Heinonen et al., 2015) |
| MX dynamin like GTPase 1 | ↑ *MX1*<sup>a</sup> | TNF-α induced cellular senescence (Kandhaya-Pillai et al., 2017) |
| MX dynamin like GTPase 2 | ↑ *MX2*<sup>a</sup> | TNF-α induced cellular senescence (Kandhaya-Pillai et al., 2017) |
| poly(ADP-ribose) polymerase family member 9 | ↑ *PARP9* | Silences pro-inflammatory genes, found in atherosclerotic plaques (Iwata et al., 2016) |
| cystatin C | ↑ *CST3*<sup>a</sup> | Atherosclerosis (Chung et al., 2018); cellular senescence (Basisty et al., 2019) |
| sterile alpha motif domain containing 9 | ↑ *SAMD9* | Anti-inflammatory factor (He et al., 2019) |
| TNF superfamily member 13b | ↑ *TNFSF13B* | Reg proliferation and differentiation of atherogenic B cells (Kyaw et al., 2013a) |
| cysteine rich secretory protein LCCL domain containing 2 | ↑ *CRISPLD2* | Regulates anti-inflammatory effects (Himes et al., 2014) |
| erythrocyte membrane protein band 4.1 like 3 | ↑ *EPB41L3* | Positively correlated smoking gene and associated with cancer (Charlesworth et al., 2010) |
| heat shock protein family A (Hsp70) member 1B | ↑ *HSPA1B* | Overexpressed in advanced atherosclerosis (Kilic and Mandal, 2012) |
| thrombospondin 1 | ↑ *THBS1* | Regulation of monocyte mobility, vascular inflammation (Liu et al., 2015) |
| interferon induced protein with tetratricopeptide repeats 5 | ↑ *IFIT5*<sup>a</sup> | Up in TNF-alpha mediated cellular senescence (Kandhaya-Pillai et al., 2017) |
| formyl peptide receptor 2 | ↑ *FPR2* | Increased in atherosclerotic lesions and plaque stability (Petri et al., 2015) |
| interferon induced with helicase C domain 1 | ↑ *IFIH1*<sup>a</sup> | Up in TNF-alpha mediated cellular senescence (Kandhaya-Pillai et al., 2017) |
| **B** |  |  |
| major histocompatibility complex, class II, DQ alpha 1 | ↓ *HLA-DQA1* | Presents peptides from extracellular protein in T cells (Zajacova et al., 2018) |

<sup>a</sup>indicates gene associated with cellular senescence

**Figure 6.**
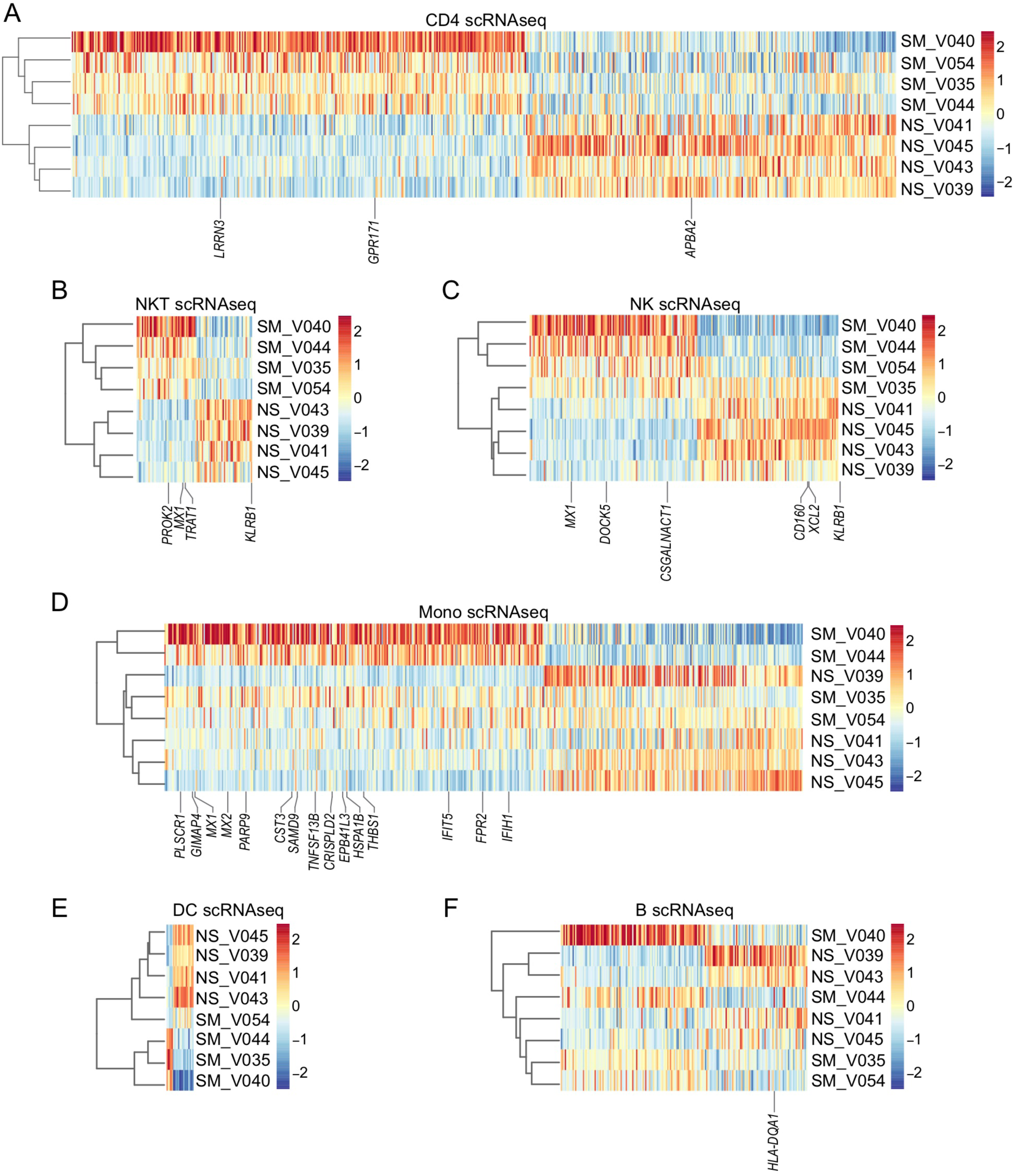
Major cell types show gene expression changes in peripheral blood of smokers. (A) Hierarchical clustering of the top 350 increased and decreased DEGs between smokers and nonsmokers from CD4 T cells (scRNAseq). (B-F) Heatmaps of smoking DE Gs (scRNAseq) in NKT cells (B), NK cells (C), monocytes (D), DCs (E), and B cells (F).

## DISCUSSION

Sharing transcriptome signatures with NK-like cells, our study reveals CD16+ CD8 T cells as elevated in smokers. This unique CD8 T subset was uncovered by scRNAseq and confirmed by mass cytometry in human PBMCs from multiple individuals. The NK-like CD8 T cells displayed downregulation of genes expressed in naïve CD8 T cells and upregulation of genes characteristic of NK cells. They shared transcriptomic features with iTNK cells, which acquire NK surface receptors and have increased cytotoxic potency (Li et al., 2010). Combined with CD16 and CD45RA protein expression, the transcriptome of CD8T-8 cells implies an innate-like, terminally-differentiated CD8 T subset. CD16, commonly associated with NK cells, acts as a receptor that binds IgG antibodies to mediate antibody-dependent cellular cytotoxicity (ADCC); exogenous or endogenous CD16 expression enables T cells to execute ADCC (Clemenceau et al., 2006; Clemenceau et al., 2008). Consistent with a heightened cytotoxic potential, the CD8T-8 cells had elevated mRNA expression of cytolytic effector molecules, *GZMB* and *PRF1*; these transcripts were also higher in smokers than nonsmokers in the NK-like CD8 T subset. Granzyme B and perforin expressing CD8 T cells contribute to the development of atherosclerotic plaques in mice (Hiebert et al., 2013; Kyaw et al., 2013b). As such, our results highlight a potential link between smoking-induced functional changes in human CD8 T cells and atherosclerosis.

As the first study to apply scRNAseq and mass cytometry to PBMCs from tobacco smoke-exposed individuals, we show that major immune populations can be discerned, and disparate subsets can be identified for CD4 T cells, CD8 T cells, NK cells, monocytes, and B cells. We did not observe any significant changes in subset distribution for NK cells, monocytes, and B cells. However, pseudobulk analysis revealed smoking DEGs, several of which were confirmed in bulk cell-type fractions. The increase in T_regs_ in smokers identified by scRNAseq data is likely masked in bulk data from isolated CD4 T cells because T_regs_ are a low frequency subset. Notably, T_regs_ have been shown to induce T-cell senescence (Ye et al., 2012), highlighting a potential role for the increase in T_regs_ observed in smokers. In addition, we identified one, two, four, and five senescence-related genes in CD4 T cells, NKT cells, NK cells, and monocytes (Table 1). *MX1*, a gene that is induced in TNFα-mediated senescence (Kandhaya-Pillai et al., 2017), increased in smokers in NKT cells, NK cells, and monocytes. Alluding to shared regulation of pro-senescent and pro-atherosclerotic signaling, TNFα-induced senescence genes are enriched for atherosclerosis signaling genes (Kandhaya-Pillai et al., 2017). *CST3*, which increased in smoker monocytes, is associated with subclinical atherosclerosis (Chung et al., 2018) and cellular senescence (Basisty et al., 2019). Among the monocyte smoking DEGs, we also identified *TNFSF13B*, a critical regulator of atherogenic B cell proliferation and differentiation (Kyaw et al., 2013a). Other notable genes connected to atherosclerosis that increased in smokers’ monocytes include *PARP9* (Iwata et al., 2016), *HSPA1B* (Kilic and Mandal, 2012), and *FPR2* (Petri et al., 2015).

We used established markers and trajectory inference to characterize the CD8 T differentiation state shifts between smokers and nonsmokers. Both approaches demonstrate that smokers lose naïve and gain T_CM_-like and T_EMRA_-like cells. Composition shifts in CD8 T cells established by scRNAseq were reflected in bulk methods. Interestingly, all three transcriptomic techniques identified gene expression changes associated with PD-1^hi^ CD8 T cells. With persistent antigen stimulation, the inhibitory effect of the PD-1 pathway contributes to pathologies associated with T-cell dysfunction during chronic viral infection and tumor evasion of host immune response (Duraiswamy et al., 2011). Duraiswamy et al. (2011) demonstrated that PD-1^hi^ CD8 T cells obtained from healthy adults had similar gene expression profiles to PD-1^lo^ CD8 T cells and did not show either exhausted gene signatures or phenotypes characteristic of PD-1^hi^ cells obtained from humans or mice with chronic infections. Smoking DEGs that were found in CD8 T cells in pseudobulk scRNAseq and confirmed in the microarray were included within these gene signatures (Figure 5H and Table 1). Notably, bulk RNAseq was run on samples from a prior visit to that of samples used for scRNAseq, suggesting a chronic or recurring state of activation in smokers’ CD8 T cells.

The NK-like CD8 T cells were found to have the latest pseudotimes, consistent with an end-stage T_EMRA_ phenotype. The loss of naïve and accumulation of terminally differentiated T cells, observed here in smokers, mimics the altered distribution of T cell subsets reported in aging and chronic infections that is proposed to result from repeated or persistent stimulation of immune cells, ultimately leading to loss of immune function either due to replicative senescence or functional exhaustion (Akbar and Henson, 2011; Larbi and Fulop, 2014; Reiser and Banerjee, 2016). Gene expression changes in low-frequency subsets may not be discernable in pseudobulk and bulk analyses. Therefore, we looked for indicators of T cell dysfunction within the smoking-associated NK-like CD8 T subset. While *TOX*, a transcription factor that controls fate commitment in exhausted T cells (Khan et al., 2019), was elevated in smokers’ NK-like CD8 T cells compared to other CD8 T cells, it was only detected in 8.9% of cells within this cluster. Other exhaustion markers *PDCD1, CTLA4*, and *HAVCR2* were not increased. Whereas exhausted CD8 T cells lack cytotoxic activity, the high expression of genes encoding proteins responsible for cytolytic activity in CD8T-8 cells suggests that these cells more likely represent a senescent or pre-senescent state. Supporting NK-like CD8 T cells as (pre-)senescent, smokers and nonsmokers had elevated expression of *KLRG1*, an inhibitory receptor correlated with extensive proliferative history (Voehringer et al., 2001), and *B3GAT1* (CD57), a marker of limited proliferative potential and shortened telomeres (Brenchley et al., 2003). Senescence can be induced as the result of telomere shortening or non-telomeric DNA damage (Akbar and Henson, 2011), both of which have been reported to occur in smokers (Song et al., 2010; Valdes et al., 2005). Accelerated immune system aging accompanies T cell senescence and can manifest as impaired immunological memory (Reading et al., 2018), which could contribute to attenuated immune responses in smokers. Of note, *GFI1*, a transcriptional repressor of IL-7Rα that drives terminal differentiation of CD8 T cells (Ligons et al., 2012), was elevated in CD8T-8 cells. *GFI1* is a reproducible methylation biomarker for tobacco smoke exposure (Bacher and Kendziorski, 2016; Joubert et al., 2016; Joubert et al., 2012; Parmar et al., 2018), and altered methylation persists up to at least 30 years after smoking cessation (Joehanes et al., 2016). Taken together, this indicates that epigenetic modifications likely contribute to smoking-mediated reprogramming of CD8 T cells. The acquisition of CD16 and NK-like characteristics would imply an underappreciated role for CD16 receptor in maintenance of cytotoxic activity in T_EMRA_ cells in smokers.

In conclusion, our discovery uncovers a new immune-cell subtype that can be isolated to investigate how NK-like CD8 T cells contribute to proinflammatory state(s) in smoking-mediated chronic inflammatory conditions. Our data illustrates novel links between smoking-induced gene expression changes and both atherosclerosis and senescence, in multiple immune populations. Consequently, our use of recently developed single-cell technologies to address tobacco smoke exposure has great potential to impact global health.

## METHODS

### Human Subjects

All donors were recruited with written informed consent under approved human IRB protocols (NIEHS 10-E-0063) by the NIEHS Clinical Research Unit between March 2013 to August 2017 from the Raleigh, Durham and Chapel Hill, NC area (Su et al., 2016; Wan et al., 2018). Whole blood was obtained from healthy (without acute disease according to self-reported medical histories) from nonsmokers, not having smoked >100 cigarettes in their lifetime, and smokers who reported their average daily cigarette consumption for the past 3 months. Serum nicotine/cotinine levels were measured by HPLC-MS (Quest, Inc.) as an indication of their smoking exposure status. Donors were recalled matching nonsmokers/smokers on age, sex and ethnicity for additional whole blood collection, cotinine levels were measured from the additional sample. See Table S1 for additional donor information.

### PBMC Isolation for scRNAseq and Mass Cytometry

Whole blood was diluted 1:5 v/v with QIAGEN Buffer EL and incubated at room temperature (RT) until clarified (∼10 min) before centrifugation (300g, RT, 10 min). After supernatant removal, leukocytes were resuspended in the same volume of Buffer EL (5 min) before spinning (300g, 8 min). Leukocytes were then washed twice in autoMACS Running Buffer (Miltenyi Biotec), counted, and cryopreserved [20% Iscove’s Modified Dulbecco’s Medium (IMDM), 70% Fetal Bovine Serum (FBS), 10% Dimethyl sulfoxide (DMSO)] at a concentration of 1×10^7^ cells/mL. Cryopreserved cells were thawed in nonsmoker/smoker pairs following the10X Genomic’s protocol for “Fresh Frozen Human Peripheral Blood Mononuclear Cells for Single Cell RNA Sequencing”. Briefly, cells were serially diluted dropwise in complete media (IMDM,10% FBS) adding 50U/mL Benzonase (Millipore Sigma) for the first dilution. After centrifugation (1100 rpm, 8 min, RT), cells were resuspended in complete media and incubated with CD15 Dynabeads (Thermo Fisher Scientific) according the manufacturer’s instructions to deplete the neutrophils from the PBMCs. PBMCs were then counted for viability and aliquoted for scRNAseq or mass cytometry in parallel.

### PBMC Preparation for Purified Cell Fractions

The mononuclear layer was isolated directly from whole blood using density gradient centrifugation with Histopaque-1077 Ficoll and ACCUSPIN™ Tubes (Sigma Millipore). Purified CD4+, CD8+, CD14+, CD19+, CD56+ cell fractions were collected using antibody-coated magnetic beads (Dynabeads,Thermo Fisher Scientific; CD56, Miltenyi Biotec). Antibody purified fractions were then extracted for DNA and RNA using the AllPrep DNA/RNA/miRNA Universal Kit according to the manufacturer’s instructions (QIAGEN).

### Mass Cytometry

Thawed PBMCs (∼3×10^6^ cells) were spun (300 g, 5 min) and resuspended in calcium magnesium-free phosphate buffered saline (PBS). 1µM Cisplatin (Fluidigm) was added for viability staining for 5 minutes before quenching the reaction with MaxPar Cell Staining Buffer (Fluidigm). After centrifugation (300 g, 5 min), cells were resuspended in CBS at a concentration of 60×10^6^ cells/mL and incubated (RT,10 min) with Fc receptor binding inhibitor before adding 25 MaxPar metal-conjugated antibodies (Fluidigm) against immunophenotypic markers for an additional 30-minute incubation at RT. Stained cells were then washed two times before resuspension in MaxPar fix and perm buffer with 125µM 191/193Ir intercalator for either an hour at RT or 4°C overnight. Cells were then washed twice with CSB and two times with Nuclease-Free water (Thermo Fisher Scientific) followed by filtering through 40uM strainers to remove aggregates. Cells were then counted and resuspended in nuclease-free water at ∼ 5×10^5^ cells/mL with 1:10 volume of four-element calibration beads (Fluidigm) and analyzed on a Helios instrument (Fluidigm) for 250,000 events for each donor at the NIEHS Flow Cytometer Center. Following the manufacturer’s instructions, downstream processing involved normalization by the calibration beads and .fcs files were uploaded to Cytobank.

Events were gated in Cytobank to identify single viable cells. Cells gated from spiked-in normalization beads were subsequently gated by Iridium (191Ir) and Cisplatin (198Pt) to obtain DNA positive cells. Single cells were identified by event length and Iridium (193Ir) and viable cells by Cisplatin-198Pt and leukocyte marker CD45. Viable cells were exported as fcs files and imported into VorteX (Samusik et al., 2016) using all events for each donor totaling 990,748 cells. Using the default parameter recommendations (Kimball et al., 2018), all data were transformed using hyperbolic arcsin (f=5). Applying a noise threshold of 1.0, clustering analysis was performed using a Euclidean length profile of 1.0 in X-shift and the weighted K-nearest neighbor density estimation (K). An elbow point validation was performed to determine the optimal cluster K value (K=25) which was then used to create a Force-Directed Layout (FDL) for visualization. 136 clusters were identified from the 990,748 events, one cluster was determined to be red blood cells (RBCs; positive expression profile for CD235a/b) and 13 clusters had multiple lineage markers and were determined to be doublets (e.g. positive expression profiles for CD19 [B cells] and CD3 [T cells]) which was a total of 6900 cells that were removed prior to downstream analysis.

### Transcriptomics

For scRNAseq, libraries were prepared with the Chromium™ Single Cell 3’ Library & Gel Bead Kit v2 (10X Genomics). For bulk RNAseq, RNA from isolated CD8+ cell fractions was prepared using the TruSeq Stranded Total RNA Library Prep Gold (Illumina).

scRNAseq data was processed with CellRanger. Uniquely aligned reads sharing equal barcode×UMI tags but annotated to multiple protein-coding transcripts (i.e. ambiguous UMIs), within each replicate were discarded from the analysis. Dataset alignment, SNN clustering, and UMAP visualization of scRNA data were performed with Seurat v3 (Stuart et al., 2019); cell cluster marker genes and smoking DEGs were identified with Seurat v2 (Butler et al., 2018) implementing the MAST algorithm (Finak et al., 2015). Slingshot was used to perform pseudotime analysis and identify temporally expressed genes (Street et al., 2018).

For microarray, differentially expressed genes were detected using log2-transformed expression fold change estimates with respect to the composite average of RMA-corrected fluorescence log-intensity levels (log2FC) across matched fractions (CD14, CD19, CD4, CD56, CD8) from multiple individual female donors, both smoking and non-smoking (N = 53 overall, with N≥5 per cell fraction × smoking status group). Probe-wise log2FC values were tested across statistical groups through a resolution-weighed ANOVA; resolution weights represented relative metrics of fluorescence discrimination in the dynamic range of detection, i.e. cumulative hazard of multivariate ANOVA significance scores (probe × cell fraction × smoking status) from probe-wise generalized linear modeling of RMA-corrected fluorescence log-intensities using an exponential distribution and inverse link function (Lozoya et al., 2018). DEGs were detected from the annotation of probes with significance level p<0.05 adjusted for multiple comparisons (Benjamini and Hochberg, 1995), then filtered against a minimum probe-wise effect size δ_log2FC_>0.3×σ_log2FC_ and post hoc pairwise significance (Student’s t-test p<0.05) between log2FC values of at least 1 matched comparison between smokers and nonsmokers on same cell fraction levels. For probe-level effect size filtering, δ_log2FC_=0.3×σ_SSR_ is 5% of the 6σ-spread log2FC regression error with respect to a probe’s grand mean [where (σ_SSR_)^2^=(SSR_log2FC_)/(N-1)] compared to 5% of the 6σ-spread in measurement error about the mean log2FC of each statistical group in the probe [where (σ_log2FC_)^2^=(SSE_log2FC_)/(N-1)].

We used GSEA (Mootha et al., 2003; Subramanian et al., 2005) to perform gene set enrichment analysis for Chemical and Genetic Perturbations and Immunological Signatures (Godec et al., 2016) gene sets for scRNAseq, bulk RNAseq, and microarray. The venn diagrams to compare DEGs among scRNAseq, bulk RNAseq, and microarray were generated with VennDiagramWeb (Lam et al., 2016).

## FUNDING DETAILS

This work was funded by the Intramural Research Program of the National Institute of Environmental Health Sciences-National Institutes of Health (Z01-ES100475) and a grant from NIH/FDA Intramural Center for Tobacco Research to D.A.B.

