## Supplemental Table 1 and Supplemental Figures for "Single-cell analyses identify tobacco smoke exposure-associated, dysfunctional CD16^+^ CD8 T cells with high cytolytic potential in peripheral blood"

**Supplemental Table 1. Donor Information**

| Sample ID[a] | Race[b] | Sex[c] | Age | BMI | Smoking Status[d] | Cotinine (ng/ml) | Cigarettes per day | Yrs Smoking | AHRR cg05575921[e] | CD8 Bulk RNAseq | Microarray |
|---|---|---|---|---|---|---|---|---|---|---|---|
| F001 | AA | F | 48 | 30.3 | SM | 292 | 50 | 36 | 0.64 | N | CD8, CD4, CD56, CD14, CD19 |
| F002 | AA | F | 39 | 30.8 | NS | <2 | | | 0.82 | N | CD8, CD4, CD56, CD14, CD19 |
| F003 | AA | F | 46 | 22 | NS | <2 | | | 0.89 | N | CD8, CD4, CD56, CD14, CD19 |
| F005 | AA | F | 53 | 32 | SM | 344 | 20 | 22 | 0.75 | N | CD8, CD4, CD56, CD14, CD19 |
| F007 | AA | F | 55 | 37.8 | SM | 410 | 20 | 41 | 0.55 | N | CD8, CD4, CD56, CD14, CD19 |
| F014 | AA | F | 53 | 67.3 | NS | <2 | | | 0.92 | N | CD8, CD4, CD56, CD14, CD19 |
| F015 | AA | F | 46 | 25.4 | NS | <2 | | | 0.90 | N | CD8, CD4, CD56, CD14, CD19 |
| F016 | AA | F | 51 | 29.8 | SM | 273 | 21 | 21 | 0.70 | N | CD8, CD4, CD56, CD14, CD19 |
| F017 | AA | F | 52 | 45.3 | NS | <2 | | | 0.89 | N | CD8, CD4, CD56, CD14, CD19 |
| F018 | AA | F | 47 | 16.1 | SM | 186 | 20 | 29 | 0.71 | N | CD8, CD4, CD56, CD14, CD19 |
| F025 | W | F | 31 | 19.2 | NS | <2 | | | 0.88 | Y | CD8, CD4, CD56, CD14, CD19 |
| F026 | W | F | 50 | 26.3 | NS | <2 | | | 0.88 | Y | CD8, CD4, CD56, CD14, CD19 |
| F031 | W | F | 41 | 29.6 | SM | 95 | 20 | 28 | 0.70 | Y | CD8, CD4, CD56, CD14, CD19 |
| F033 | W | F | 47 | 26.6 | NS | <2 | | | 0.89 | Y | CD8, CD4, CD56, CD14, CD19 |
| F037 | W | F | 45 | 21.1 | SM | 182 | 12 | 19 | 0.61 | Y | CD8, CD4, CD56, CD14, CD19 |
| F040 | W | F | 55 | 23.3 | SM | 210 | 20 | 41 | 0.54 | Y | CD8, CD4, CD56, CD14, CD19 |
| F043 | W | F | 45 | 28.7 | NS | <2 | | | 0.88 | Y | CD8, CD4, CD56, CD14, CD19 |
| F047 | W | F | 48 | 34.1 | NS | <2 | | | 0.88 | Y | CD8, CD4, CD56, CD14, CD19 |
| F048 | W | F | 47 | 25.5 | SM | 122 | 20 | 29 | 0.59 | N | CD8, CD4, CD56, CD14, CD19 |
| F054 | AA | F | 43 | 31.2 | SM | 147 | 20 | 28 | 0.57[f] | N | CD56 |
| F061 | W | F | 32 | 18.7 | SM | 111 | 10 | 18 | 0.49 | Y | |
| F067 | AA | F | 48 | 30.7 | SM | 157 | 10 | 10 | 0.50[f] | N | CD56 |
| F075 | W | F | 25 | 25.3 | NS | <2 | | | 0.87 | Y | |
| F093 | W | M | 42 | 28.7 | NS | <2 | | | 0.90[f] | Y | |
| F120 | W | F | 30 | 24.4 | NS | <2 | | | 0.91[f] | Y | |
| F160 | W | F | 47 | 27.3 | SM | 143 | 15 | 32 | 0.71[g] | N | CD56 |
| F166 | W | F | 51 | 28.7 | NS | <2 | | | 0.86[g] | N | CD56 |
| F198 | W | F | 51 | 34.4 | SM | 327 | 20 | 20 | 0.59[f] | Y | |
| F209 | W | M | 55 | 20.7 | SM | 269 | 18 | 35 | 0.52[f] | Y | |
| F239 | W | F | 48 | 33.2 | SM | 215 | 9 | 33 | 0.89[f] | Y | |
| V035(F031) | W | F | 43 | 30.1 | SM | 511 | 25 | 30 | See F031 | N | |
| V039(F093) | W | M | 44 | 29.6 | NS | <2 | | | See F093 | N | |
| V040(F198) | W | F | 52 | 34.0 | SM | 358 | 20 | 21 | See F198 | N | |
| V041(F043) | W | F | 47 | 27.0 | NS | <2 | | | See F043 | N | |
| V043(F047) | W | F | 50 | 32.9 | NS | <2 | | | See F047 | N | |
| V044(F037) | W | F | 48 | 20.4 | SM | 240 | 10 | 22 | See F037 | N | |
| V045(F120) | W | F | 31 | 22.1 | NS | <2 | | | See F120 | N | |
| V054(F209) | W | M | 56 | 20.7 | SM | 357 | 18 | 36 | See F209 | N | |

[a]Sample ID with V correspond to second visit from individual with sample ID beginning with F. [b]AA=African American; W= White. [c]F=Female; M=Male.
[d]SM=Smoker; NS=Nonsmoker. [e]450K methylation or [f]850K in Whole Blood or [g]CD56 fraction.

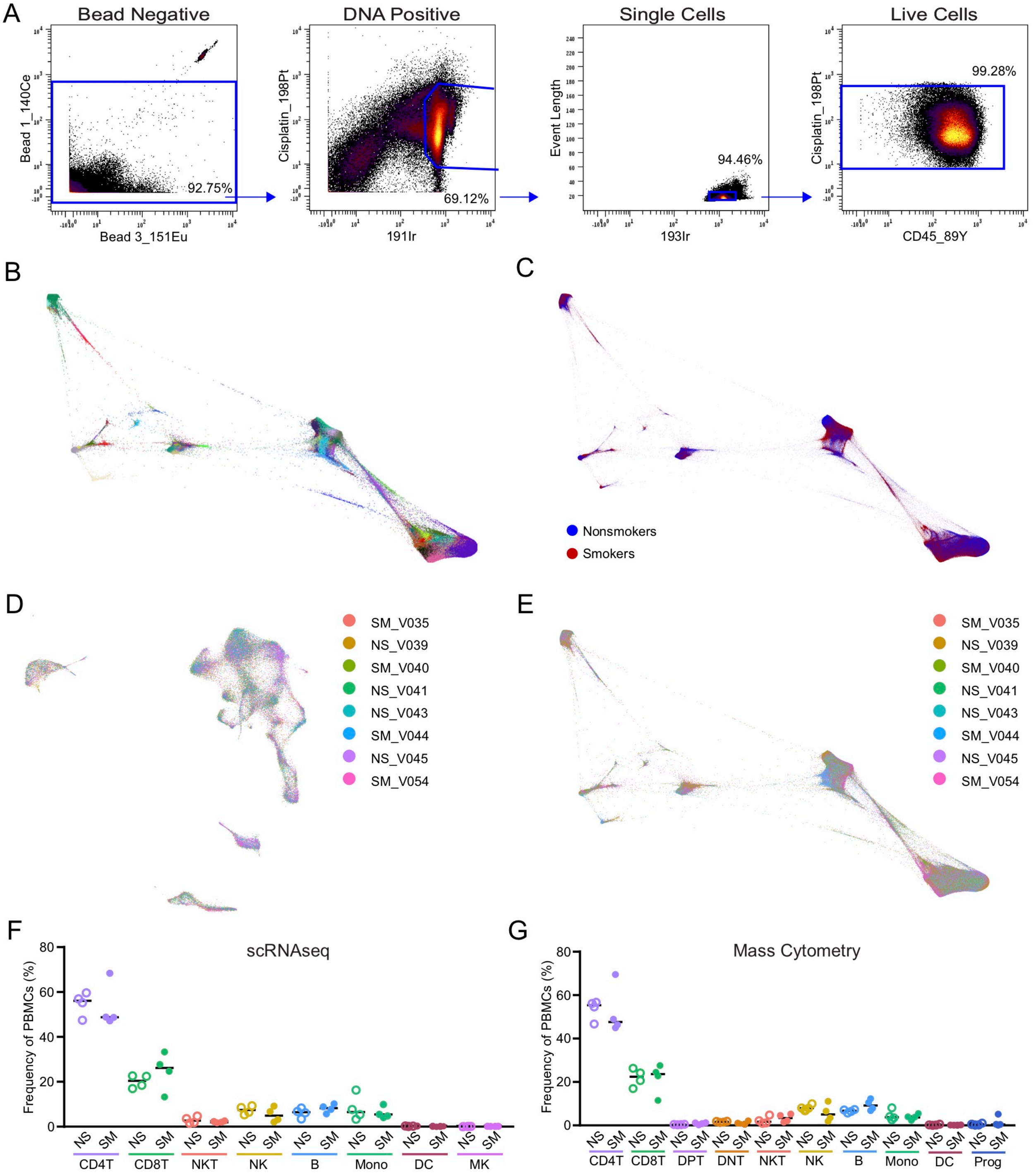


**Figure S1. ScRNAseq and mass cytometry profiling of PBMCs, Related to Figure 1**

(A) Mass cytometry biaxial plots of the gating strategy. Cells gated from spiked-in normalization beads are subsequently gated by Iridium (191Ir) and Cisplatin (198Pt) to obtain DNA positive cells. Single cells are identified by event length and Iridium (193Ir) and viable cells by Cisplatin-198Pt and leukocyte marker CD45.

(B-C) FDL mass cytometry of 122 immune cell cluster IDs displayed by cluster ID color (B) and smoking status (C; nonsmokers blue, smokers red).

(D) scRNAseq UMAP as described in Figure 1B. Cells are colored by individual donors.

(E) Mass cytometry FDL as described in Figure 1C. Cells are colored by individual donors.

(F-G) Frequencies of PBMCs by major cells types compared between smokers (filled) and nonsmokers (unfilled) for scRNAseq (F) or mass cytometry (G) did not show significant differences. Bar = median, Mann-Whitney U test.

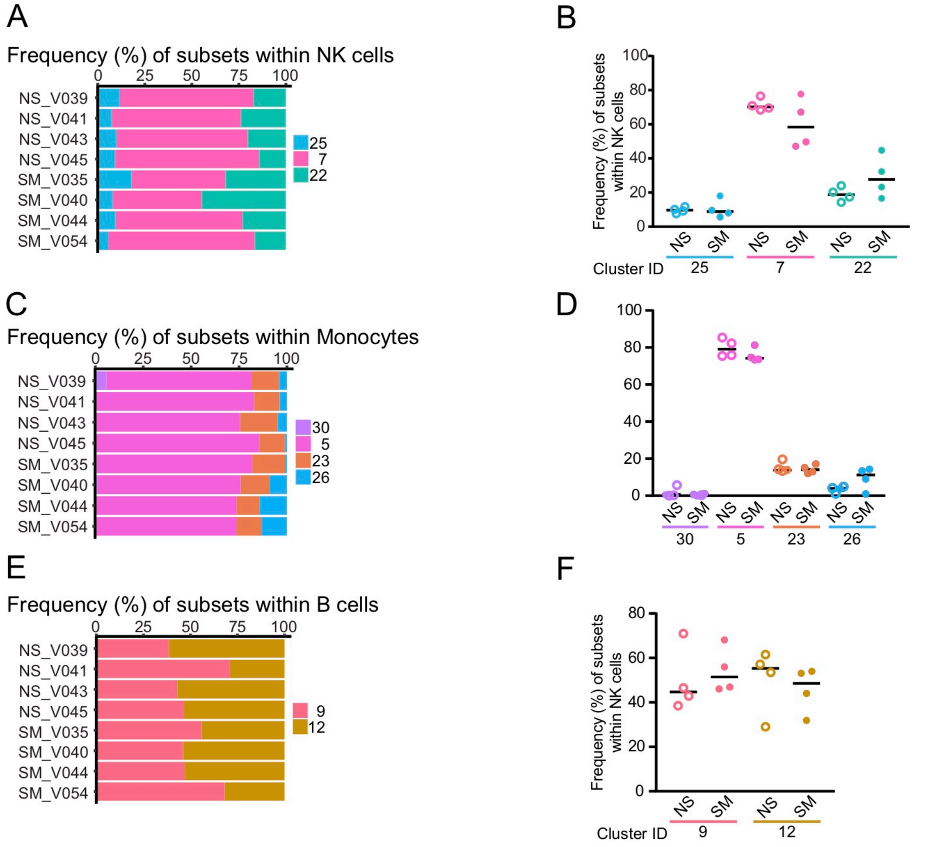


**Figure S2. Subtype distributions within other major cell populations do not shift between smokers (SM) and nonsmokers (NS), Related to Figure 2**

(A-B) Frequency of subsets within NK cells. Individual donor distribution of NK cell subsets (A). Smokers (filled) had no significant shifts in NK cell frequency compared to nonsmokers (unfilled). Bar = median, Mann-Whitney U test (B).

(C-D) Frequency of subsets within monocytes. Individual donor distribution of monocyte subsets (C). Smokers (filled) had no significant shifts in monocyte frequency compared to nonsmokers (unfilled). Bar = median, Mann-Whitney U test (D).

(E-F) Frequency of subsets within B cells. Individual donor distribution of B cell subsets (E). Smokers (filled) had no significant shifts in B cell frequency compared to nonsmokers (unfilled). Bar = median, Mann-Whitney U test (F).

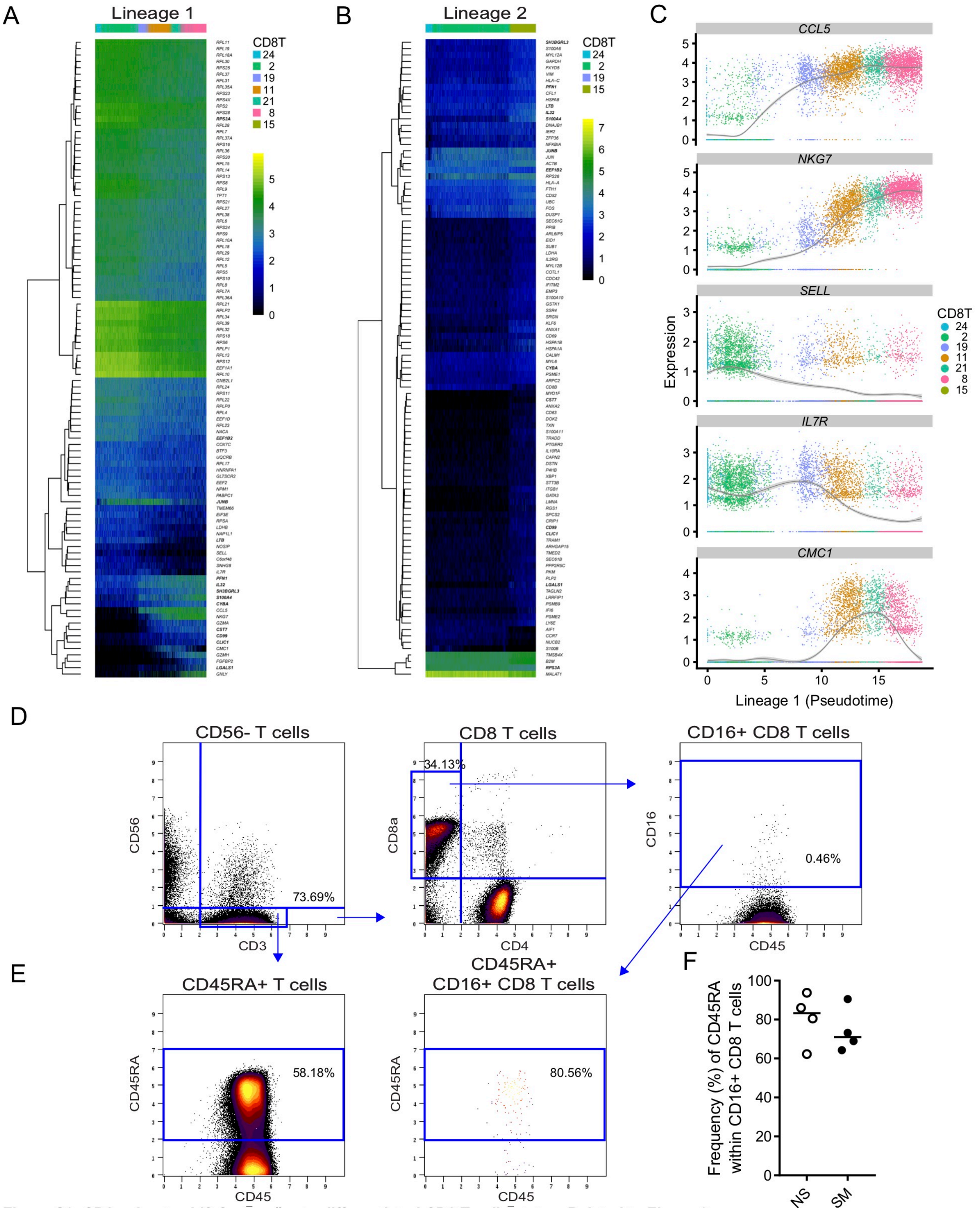


**Figure S3. CD8 subsets shift from naïve to differentiated CD8 T cells states, Related to Figure 3**

(A-B) Pseudotemporal heatmaps of CD8 T cell differentiation of lineage 1 (A) and lineage 2 (B).

(C) Pseudotemporal trajectory of CD8 T cell differentiation in Lineage 1 of CCL5, NKG7, SELL, IL7R and CMC1.

(D) Mass cytometry gating strategy to determine frequency of CD16+ CD8 T cells. Viable cells (Figure S1A, right panel) displayed in a CD3/CD56 surface marker biaxial plot. CD56 negative cells (lower right quadrant) were gated by CD4 and CD8 to obtain single positive CD8 T cells (upper left quadrant), which were then used to determine the frequency of CD16+ CD8 T cells.

(E) CD3+ T cells gated by a CD45RA/CD45 biaxial plot to establish an accurate CD45RA+ gate that was then applied to the CD16+ CD8 T cells showed the majority of cells were positive for CD45RA.

(F) Frequency of CD45RA+ cells within CD16+ CD8 T cells from smokers and nonsmokers. Bar = median, Mann-Whitney U test.

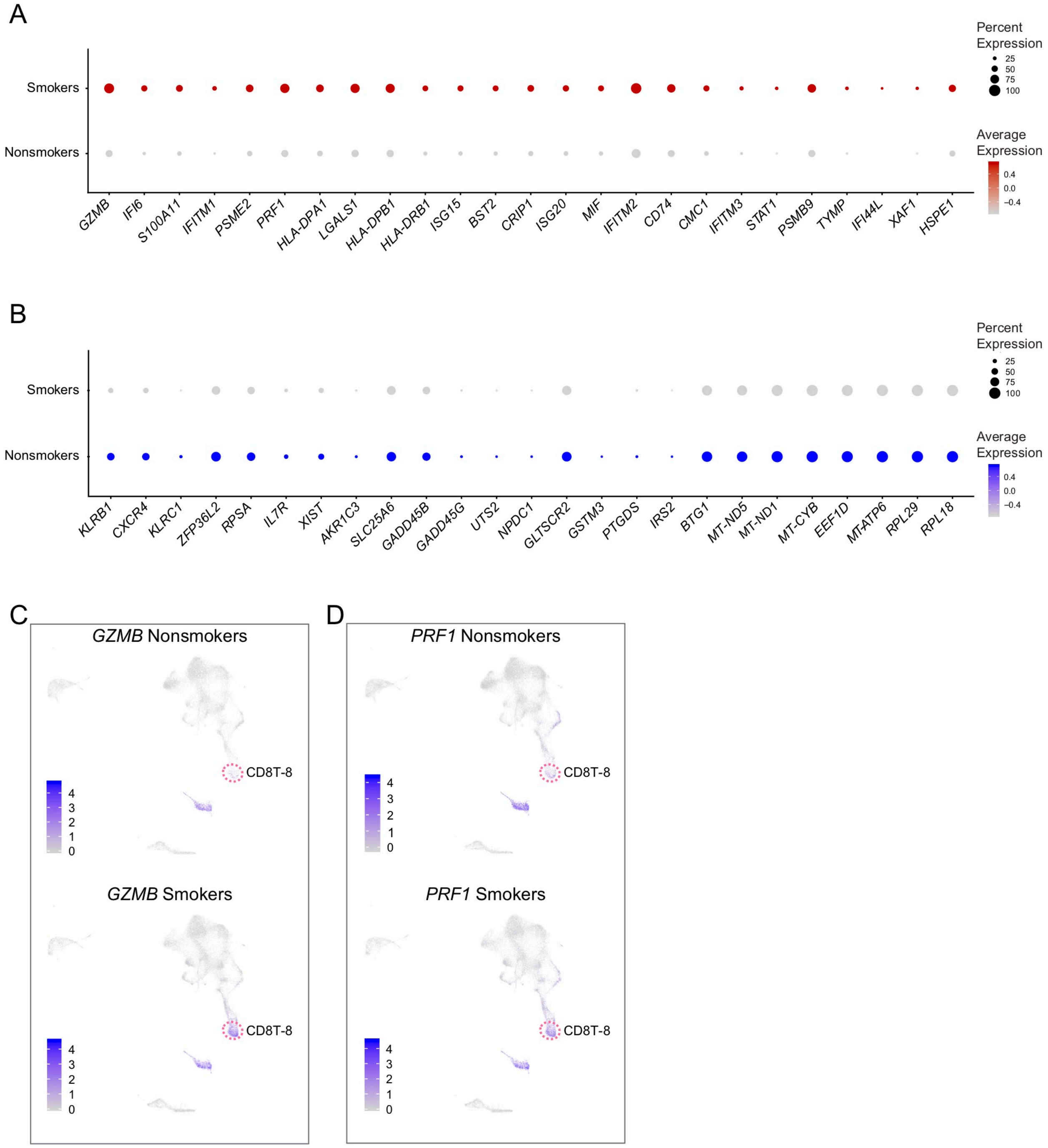


**Figure S4. CD16+ CD8 T cells characterization by scRNAseq, Related to Figure 4**

(A-B) Genes altered in smokers within the CD8T-8 subset. 25 genes with increased (A) or decreased (B) per cell gene expression were ordered by the difference in percentage of CD8T-8 cells expressing each gene between smokers and nonsmokers. Color intensity indicates average per cell expression and circle size represents the percent of cells expressing the gene.

(C-D) UMAP comparison of nonsmokers and smokers, as described in Figure 4C and 4E, displaying the increase in *GZMB* (C) and *PRF1* (D) from smokers in the CD8T-8 cluster.

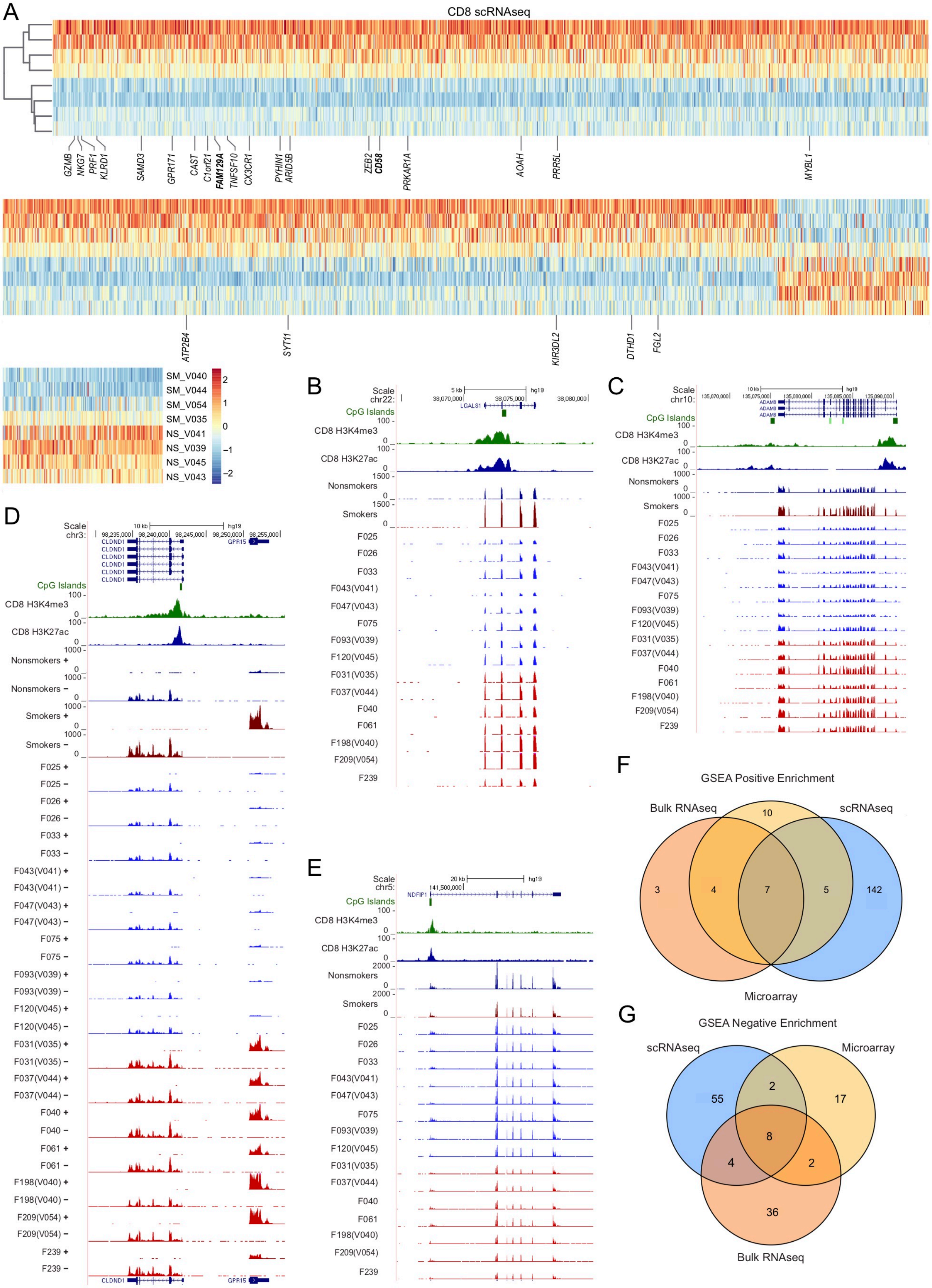


*(legend on next page)*

**Figure S5. Activation of CD8 T cells in peripheral blood of smokers, Related to Figure 5**

(A) scRNAseq heatmap of all differentially expressed genes (DEGs) between smokers and nonsmokers from seven CD8 T cells clusters. Individual donors were separated by smoking status using smoking scRNA-DEGs for hierarchical clustering. Genes labeled were also found to be significantly upregulated in the CD8 T cell microarray results (See Figure 5D).

(B-E) Genome browser tracks of DEGs from bulk RNAseq. LGALS1 (B), ADAM8 (C), and CLDND1 (D) were significantly increased in bulk RNAseq and scRNAseq data, while GPR15 (D) was increased in the RNAseq and microarray data. NDFIP1 (E) was significantly decreased in the bulk RNAseq and scRNAseq data.

(F-G) Venn diagrams of positively (F) and negatively (G) enriched gene signatures for scRNAseq, microarray and bulk RNAseq.

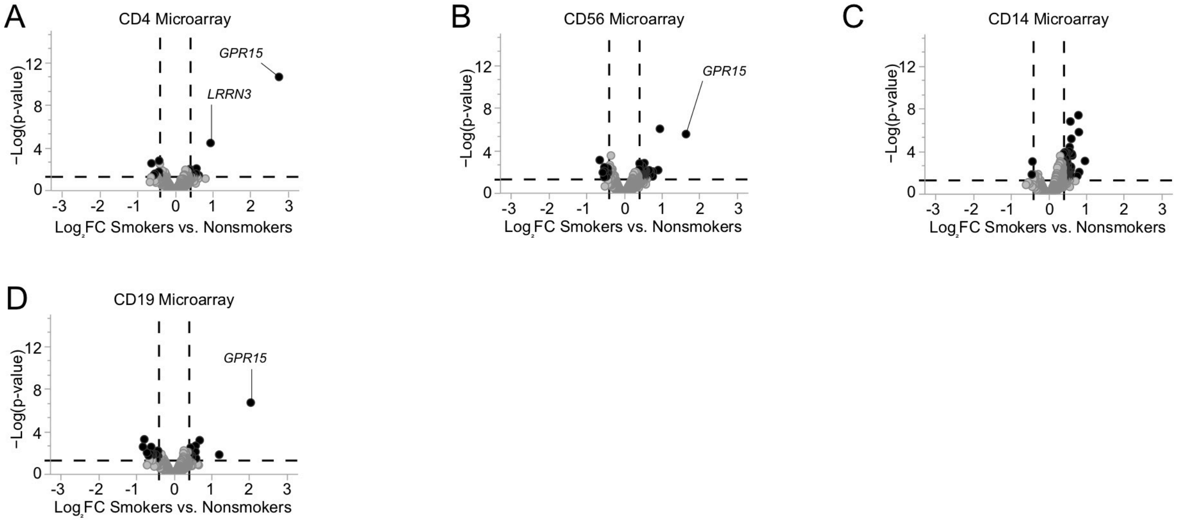


**Figure S6. Other major cell type expression changes in peripheral blood of smokers, Related to Figure 6**
(A-D) Microarray volcano plots of isolated CD4+ T cells (A), CD56+ cells (B), CD14+ cells (C) and CD19+ B cells (D).
